# Beyond energy balance in agrivoltaic food production: Emergent crop traits from wavelength-selective solar cells

**DOI:** 10.1101/2022.03.10.482833

**Authors:** Melodi Charles, Brianne Edwards, Eshwar Ravishankar, John Calero, Reece Henry, Jeromy Rech, Carole Saravitz, Wei You, Harald Ade, Brendan O’Connor, Heike Sederoff

**Affiliations:** Department of Plant and Microbial Biology, North Carolina State University, Raleigh, North Carolina, United States of America; Department of Mechanical and Aerospace Engineering and Organic and Carbon Electronics Laboratories, North Carolina State University, Raleigh, North Carolina, United States of America; Department of Physics and Organic and Carbon Electronics Laboratories, North Carolina State University, Raleigh, North Carolina, United States of America; Department of Chemistry, University of North Carolina, Chapel Hill, North Carolina, United States of America

## Abstract

The integration of semi-transparent organic solar cells (ST-OSCs) in greenhouses offers new agrivoltaic opportunities to meet the growing demands for sustainable food production. The tailored absorption/transmission spectra of ST-OSCs impacts the power generated as well as crop growth, development and responses to the biotic and abiotic environments. We grew lettuce and tomato, traditional greenhouse crops, under three ST-OSC filters that create different light spectra. Lettuce yield and early tomato development are not negatively affected by the modified light environment. Our genomic analysis reveals that lettuce production exhibits beneficial traits involving nutrient content and nitrogen utilization while select ST-OSCs impact regulation of flowering initiation in tomato. ST-OSCs integrated into greenhouses are not only a promising technology for energy-neutral, sustainable and climate-change protected crop production, but can deliver benefits beyond energy considerations.

## Introduction

Greenhouses enable the production of food crops and ornamental plants year-round outside of their natural growth zones and can therefore produce more crops on less land than conventional field cultivation, giving them an important role in feeding the world as space becomes increasingly limited. Current predictions for global food demand estimate a 70% increase by 2050, while the world population is predicted to grow by roughly 40% over the same period (1). Overall productivity in greenhouses is several times higher than in fields. A comparative analysis of tomato production showed that greenhouse productivity in New York was 12-fold higher than that of Florida fields (2), while lettuce production was more than ten times higher under greenhouse conditions on a per area basis (3).

In addition to increased productivity, greenhouses also use less water than conventional farming (3-5). This will become increasingly advantageous, because large regions of the world are expected to experience field crop losses as climate change continues to limit the availability of water for irrigation (6). Greenhouse cultivation also allows for the reduction of the ecological impacts of pesticides and fertilizers as well as a reduction in herbicide use. The enclosed nature of greenhouses presents an opportunity to more easily control and monitor chemical contaminants, such as pesticides and fertilizers, in water and soil exiting the system (3). Furthermore, greenhouses can be designed to recycle nutrients in their irrigation systems to avoid excess fertilizer use (7). This becomes especially important as the production of N-fertilizer is not only energy intensive, but its field applications lead to eutrophication of aquatic systems. Along with greater control within the enclosed environment, greenhouses shelter crops from extreme weather conditions like drought, heat or flooding, which are worsened by climate change. In 2021 alone, damage to field crops due to extreme weather events exceeded 8 billion USD (8).

However, this seemingly ideal system for plant productivity requires large amounts of energy for temperature control and supplemental lighting, which reduces its economic and environmental sustainability. Life Cycle Assessment (LCA) studies have shown that the carbon footprint of greenhouse-grown crops exceeds that of conventional crops when fossil fuels are used for heating and cooling (4, 5). The studies mentioned above that reported large improvements in greenhouse tomato and lettuce production also found energy demands nearly 19 times higher than field cultivation for tomato (2) and more than 80 times higher for lettuce (3). This prevents conventional greenhouses from being a truly sustainable method of food production. Therefore, new technologies are needed to solve the problem of high energy demand in these climate-controlled systems.

One alternative is to grow crops in insulated, fully enclosed environments that avoid the greenhouse effect caused by natural sunlight. These sole-source container farms are an alternative to greenhouse cultivation with a lower cost for climate control, but they suffer from high energy costs for artificial lighting that prevent their economic viability for crops other than microgreens or lettuce (9). These container systems can also produce crops at higher yields than conventional field agriculture (10). While these systems require less energy for space conditioning than a greenhouse (11), the elimination of natural lighting requires artificial light sources. This artificial lighting, typically provided by LEDs, is the major energy requirement and limitation of container systems (12). Indeed, one study found that such systems are only economically viable for the production of low-light crops, such as microgreens and, to a lesser extent, lettuce (9), although the high energy cost can possibly be reduced through an intermittent lighting strategy (13). Container systems trade one high energy requirement for another by excluding sunlight and relying on LEDs. The sustainable production of the majority of crops will require both the integration of a renewable energy source and better utilization of the natural sunlight available for crop production in most of the world.

While other solar-powered greenhouses do improve the sustainability of food production, they require either additional land in the form of solar farms or a reduction in yield in the case of opaque rooftop solar cells (14). Wavelength-selective semi-transparent organic solar cells (OSCs) are an alternative to both container and conventional solar power systems. The cost of OSC-greenhouses can be kept close to the cost of conventional and photovoltaic-adjacent greenhouse systems without additional land in favorable climates (15). Additionally, these OSC systems allow more light to reach the crops below, avoiding the yield losses seen in opaque systems. However, the light reaching the plants is altered in both light intensity (i.e., quantity) and light spectrum (i.e., quality). Through the selection of distinct organic semiconductors as the OSC active layer, the spectral transmission can be controlled (16). These spectrally flexible net zero energy greenhouses have been shown to be economically feasible for crop production in most regions of the world (17).

Plants depend on light for their energy supplied by photosynthesis. Light harvesting antenna complexes contain large numbers of chlorophylls that absorb primarily in the blue (400-500 nm) and red (600-700 nm) regions of the light spectrum to produce energy (18). As a result of these peak absorbance wavelengths, red and blue light are used more efficiently than green light during photosynthesis (Figure 1). The wavelengths from (400-700 nm), termed photosynthetically active radiation (PAR), are the most relevant for plant growth and are measured in PPFD, photosynthetic photon flux density. TPFD, or total photon flux density, is also used to account for the full range of wavelengths sensed by plants (19-21).

**Figure 1.**
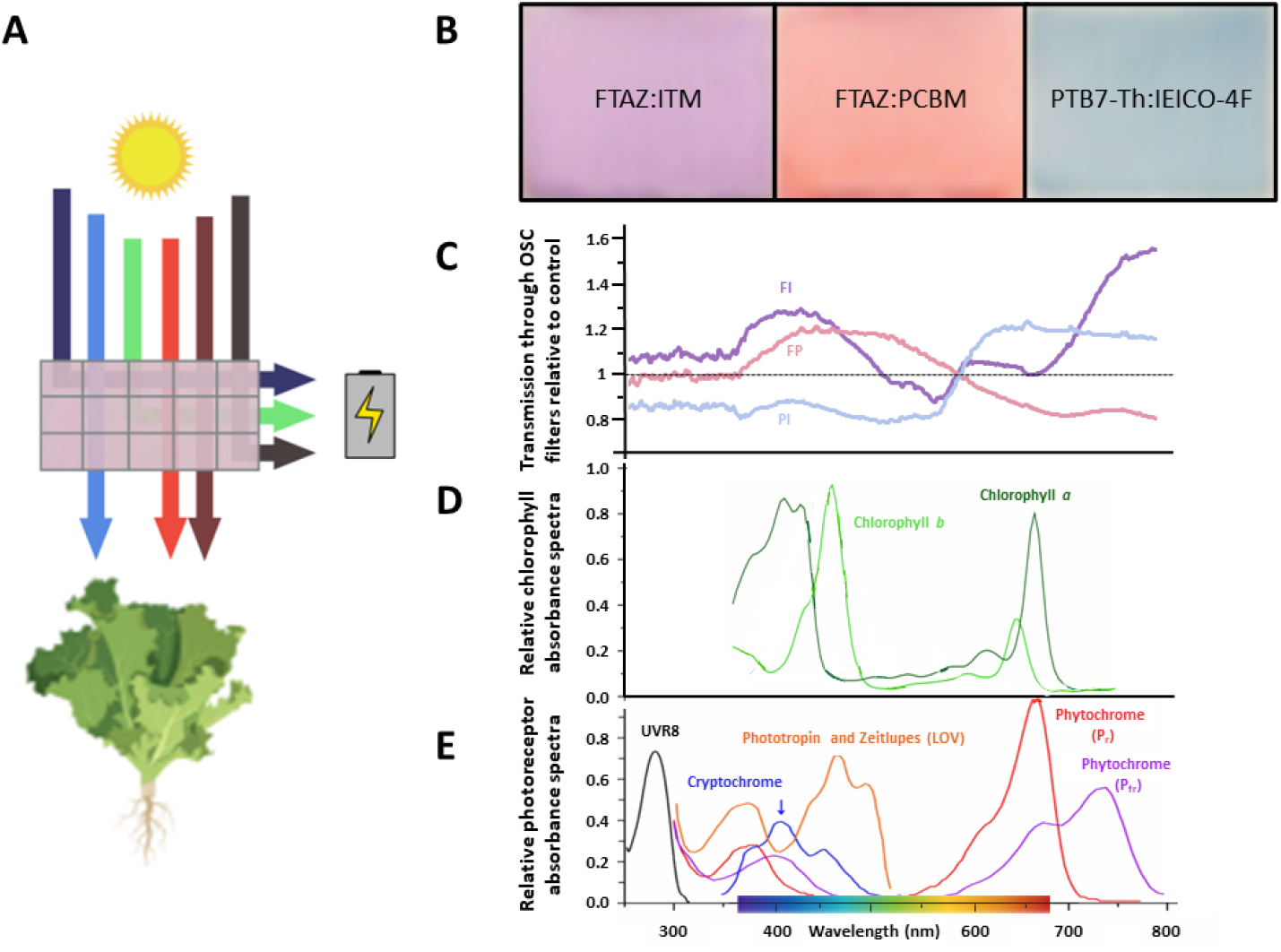
Wavelength selectivity of OSC devices. **A**) Schematic of a wavelength-selective OSC device. Specific wavelengths of light are harvested to generate electrical energy to power the greenhouse. The wavelengths most important for photosynthesis (blue, red and far-red) are selectively transmitted through the device to grow greenhouse crops. **B**) The OSC filters are named for the two organic molecules that determine their color and transmission spectrum. Different donor and acceptor molecules can be chosen to tune the wavelengths that reach the plants below. **C**) Ratio of light transmission through the OSC filters relative to the light transmitted through the control. Each filter varies in relation to the control and to each other over PAR and in the far-red region (700-750 nm). **D**) Absorption spectra of plant chlorophylls. Chlorophyll a and b harvest light energy from sunlight to power photosynthesis over PAR (400-700 nm), especially in the blue and red regions (adapted from Reference 18). **E**) Absorption spectra of photoreceptors. UVR8 absorbs UV light. Cryptochromes, phototropins and zeitlupes absorb primarily blue light. Phytochromes absorb red and far-red light (adapted from Reference 18).

Because light is the only energy source for plants, their entire growth and development is dependent upon and regulated by changes in their light environment. To sense these light changes, plants have evolved a complex network of additional chromophores called photoreceptors that sense different wavelengths, intensity, and direction in the 400-700 nm range of PAR and beyond (22). This sensory input informs photomorphogenesis, the adjustment of growth and development to light conditions (22). Photomorphogenesis occurs through large integrated gene networks with other environmental factors such as nutrient availability and temperature (23-25). Many crop production characteristics are influenced by these gene networks, including height, nutrient content and the timing of flowering and fruit production (26-28).

Due to the tunable nature of OSCs, net zero greenhouses can create a wide variety of light environments for crops. Various models have estimated that sufficient crop and energy harvests can be produced using different organic semiconductors in many climates (29, 30). Peppers (31) and tomatoes (32) have been grown in greenhouses partially covered in OSC panels with minimal impacts on growth. Flowering species, such as tomato, are important greenhouse crops that are particularly sensitive to changes in light spectrum. To address the question of how the altered spectra produced by OSC filters might impact light-sensitive growth and development of common greenhouse crops, we grew a low light-requiring crop, lettuce (*Lactuca sativa cv*. ‘Red oak leaf’) and a high light-requiring crop, tomato (*Solanum lycopersicum cv*. ‘Moneymaker’) under simulated OSC greenhouse conditions. Previously studied OSC filters were selected based on their spectral complementarity with chlorophyll and photoreceptor absorbance and ease of fabrication: FTAZ:IT-M (FI), FTAZ:PCBM (FP) and PTB7-Th:IEICO-4F (PI) (16, 33). We found no detrimental impacts on biomass accumulation in either species. Furthermore, we demonstrate modifications in gene expression detected through a transcriptome analysis that point to agronomically important emerging traits in crop physiology in response to the altered light spectra, including flowering time, nitrogen use, and nutritional content. This analysis demonstrates that OSC greenhouses may present synergistic opportunities to achieve environmentally sustainable agriculture and drive favorable crop traits.

## Results

Our experiments were designed to identify the molecular and physiological responses of lettuce and tomato plants to altered light spectra provided by different ST-OSCs. Here, we present the results of plants grown under different OSC filters with the same input of total photosynthetic active radiation. A second set of supplemental experiments was conducted where light intensity was allowed to vary as well as light spectra, based on the inherent variable transmittance of the OSC filters. Because many traits are not apparent unless directly tested (e.g., nitrogen use efficiency or drought responses), we used transcriptome analyses to identify differentially regulated genome networks as indicators for underlying changes in plant physiology and development. The results of the variable light intensity are included in the supplemental information.

### Light conditions under OSCs

Growth containers with integrated OSC filters were designed to house plants within a climate-controlled growth chamber equipped with lights that approximated the spectrum of natural sunlight (Figure S1). The distance from the light source to the top of each growth box was kept constant to simulate the irradiation on the roof of actual greenhouses in a previous experiment (33). The distance from the light source and height of the growth boxes was adjusted to minimize differences between light intensity (PPFD) between each filter treatment and control (Table 1).

**Table 1.** Light intensity and spectra under OSC filters Breakdown of the photon flux (μmol m^-2^ s^-1^) reaching the plants by color measured at the end of the experiments. OSC filters are referred to by the first letter of the acceptor and donor molecules: FTAZ:IT-M (FI), FTAZ:PCBM (FP) and PTB7-Th:IEICO-4F (PI). The percentages of each color relative to the total photon flux density (TPFD) were consistent for each filter in both experiments. TPFD was more consistent between treatments in the tomato experiment.

| Species | Lettuce |  |  |  | Tomato |  |  |  |
| --- | --- | --- | --- | --- | --- | --- | --- | --- |
| Treatment | C | FI | FP | PI | C | FI | FP | PI |
| Blue (400-500nm)<br>(% of TFPD) | 36<br>(14%) | 52<br>(15%) | 31<br>(12%) | 47<br>(16%) | 37<br>(14%) | 45<br>(16%) | 33<br>(12%) | 48<br>(18%) |
| Green (500-600nm)<br>(% of TFPD) | 76<br>(30%) | 91<br>(27%) | 64<br>(25%) | 95<br>(32%) | 76<br>(30%) | 83<br>(30%) | 74<br>(27%) | 90<br>(35%) |
| Red (600-700nm)<br>(% of TFPD) | 121<br>(48%) | 169<br>(49%) | 138<br>(54%) | 136<br>(45%) | 121<br>(48%) | 123<br>(45%) | 146<br>(53%) | 107<br>(41%) |
| Far-red (700-750nm)<br>(% of TFPD) | 20<br>(8%) | 30<br>(9%) | 23<br>(9%) | 21<br>(7%) | 46<br>(8%) | 51<br>(8%) | 55<br>(9%) | 36<br>(6%) |
| PPFD (400-700nm) | 233 | 312 | 233 | 279 | 254 | 272 | 277 | 260 |
| TFPD (400-750 nm) | 253 | 342 | 256 | 300 | 300 | 323 | 332 | 296 |

### Biomass was largely unaffected by OSC filters

Variation in lettuce and tomato growth was largely insignificant when only spectra varied. This was unsurprising given that there were few significant differences between filter treatments and controls when both light quality and intensity were altered by OSC filters (33). There were no significant differences in lettuce fresh weight, dry weight or leaf area at harvest stage (Figure 2a-d) and the transplant stage (Figure S2). The spectral differences between the three OSC filters alone were evidently not large enough to have a significant effect on biomass in lettuce.

**Figure 2.**
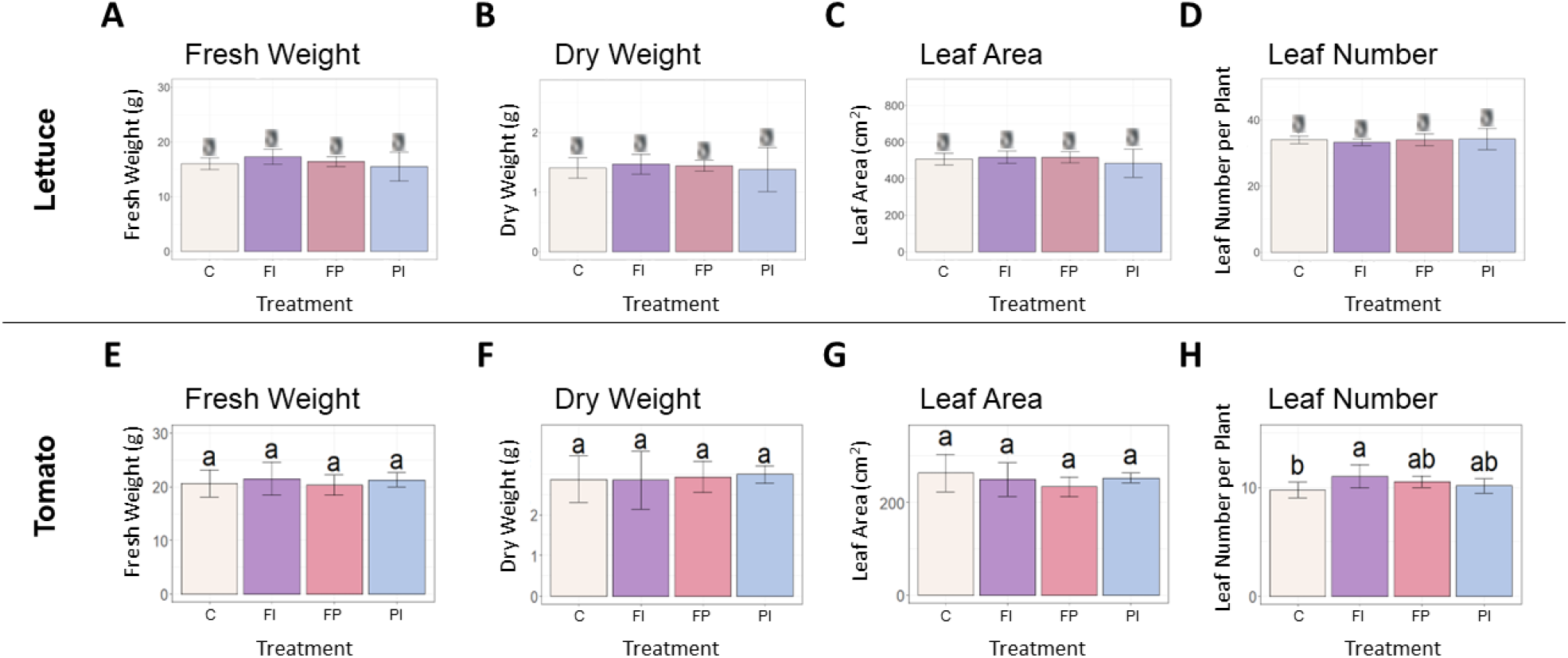
Biomass accumulation of lettuce and tomato under OSC filters. **A**) Lettuce fresh weight of lettuce grown under OSC filters and clear glass control (C). **B**) Lettuce dry weight grown under OSC filters. **C**) Lettuce leaf area under OSC filters. **D**) Number of leaves per lettuce plant under OSC filters. e-h) Tomato fresh weight, dry weight, leaf area and leaf number. Statistical significance was assessed by ANOVA and Tukey test (p<0.05). For each plot, the differences between the means of treatments marked with the same letter are not statistically significant.

Tomato biomass accumulation was measured by the same parameters recorded previously for lettuce in addition to those measurements more relevant to growth and development of a fruit crop species. As seen in lettuce, there were no significant differences in shoot fresh or dry weight between the filter treatments and control (Figure 2e, f). The FI treatment produced significantly more leaves than the control in tomato (Figure 2h). This differs from the response of lettuce to the OSC filters, where there were no significant differences between the filter treatments and the control in leaf number or other measured biomass parameters.

### Crop-specific responses in photosynthetic CO_2_-fixation and transpiration

Although overall biomass and growth phenotype did not vary significantly in lettuce plants grown under the different spectra, the rate of photosynthesis was significantly improved in the FI treatment, relative to control (Figure 3). In contrast, the transpiration rate was unaffected. Because differences in light intensity were minimized, the differences in photosynthesis are likely the result of the spectrum of the FI filter. While filter degradation did result in some variation in light intensity by the end of the lettuce experiment (Table 1, S3; Figure S3), the trends reported here remained after normalization to PPFD recorded at the time of measurement, although they were not always statistically significant (Figure S4).

**Figure 3.**
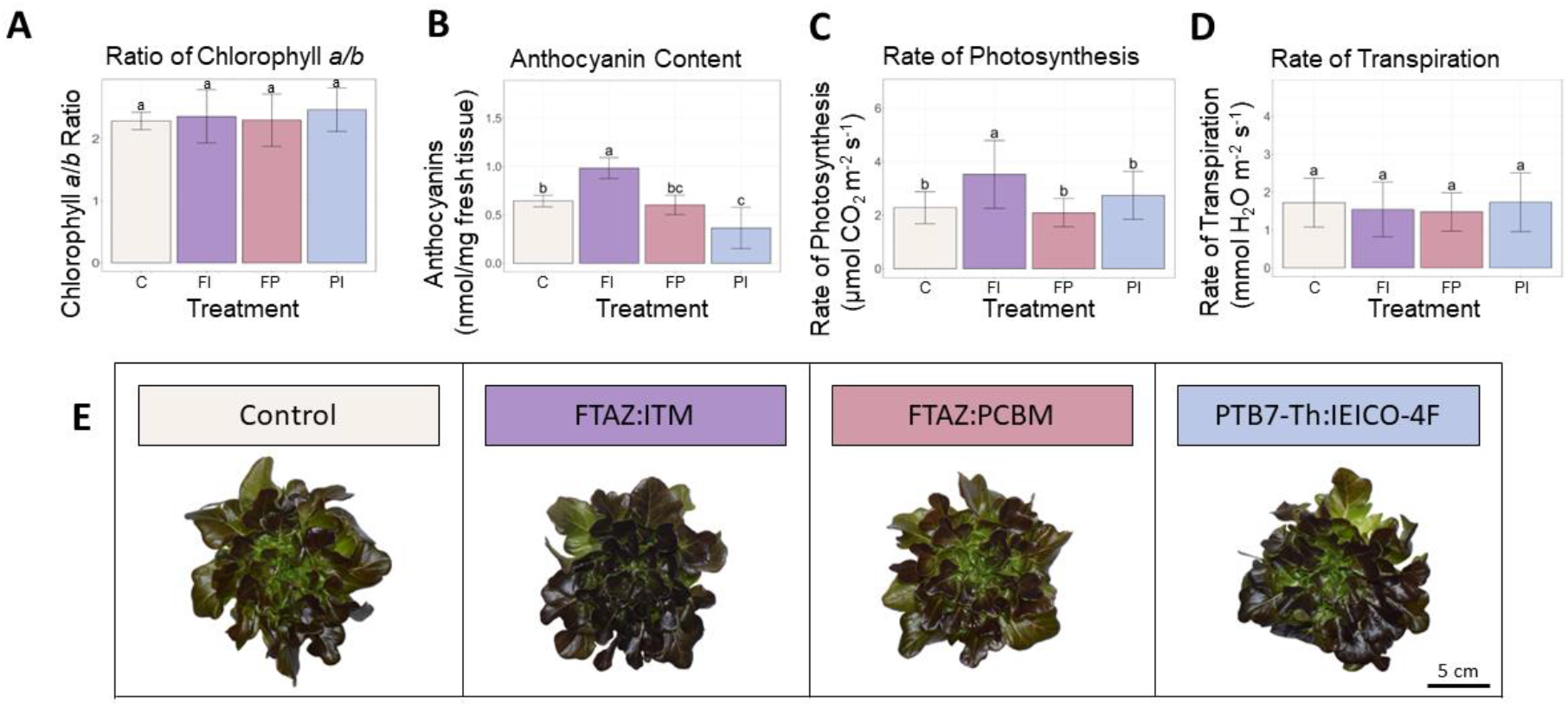
Lettuce secondary metabolites and carbon assimilation under OSC filters. **A**) Ratio of chlorophyll *a* to chlorophyll *b* in lettuce leaf tissue. **B**) Carotenoid concentration in lettuce leaf. **C**) Anthocyanin concentration in lettuce leaf tissue. c) Rate of photosynthesis (μmol CO_2_ m^-2^ s^-1^) measured under each light condition. **D**) Transpiration rate (mmol H_2_O m^-2^ s^-1^) measured under each light condition. **E**) Representative photos of lettuce plants from each treatment. Error bars represent the standard deviation. Statistical significance was assessed by ANOVA and Tukey test (p<0.05). For each plot, the differences between the means of treatments marked with the same letter are not statistically significant.

In contrast to lettuce, the tomato plants grown under the FI filter showed no significant difference in photosynthesis between control and FI filter treatment, albeit with large variation (Figure 4, S5). A significant decrease was found in the transpiration rate of these tomato plants. The increased photosynthetic rate under the FI filter in lettuce may be due to the proportion of red to blue light in relation to the amount of less efficient green light. The opposite response seen in tomato could be due to the 3% increase in green light seen in the FI tomato treatment relative to the FI treatment in the lettuce experiment. However, this increase only raises the percent of green light in the FI treatment to be equal to the percent of green in the C treatment (30%). This suggests that this response is more likely to be a species difference between lettuce and tomato, possibly affected by the sensitivity of stomatal aperture.

**Figure 4.**
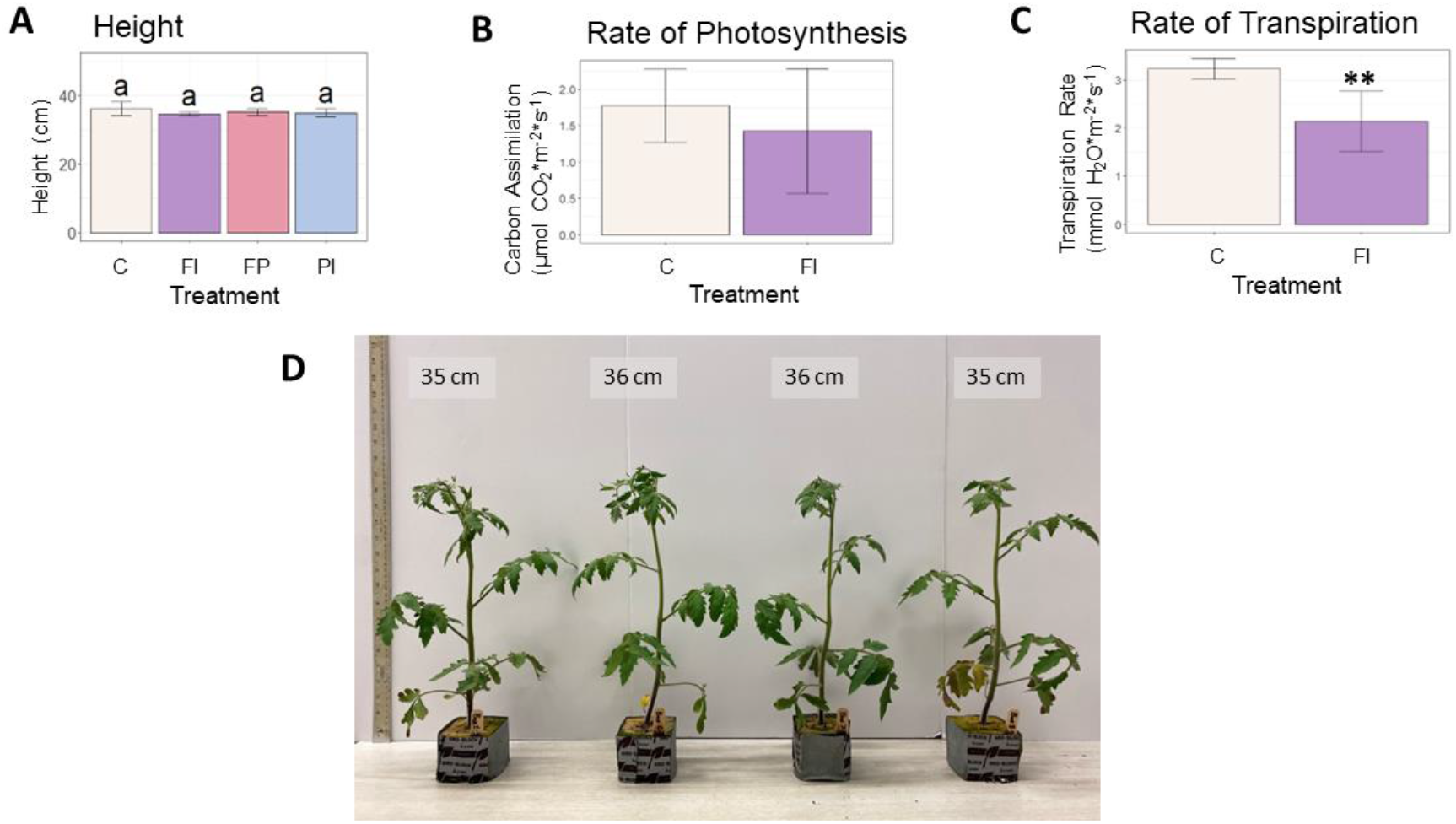
Height and carbon assimilation in tomato plants. **A**) Height of tomato plants at 30 days. **B**) Rate of photosynthesis of tomato plants under the FI filter relative to the unfiltered control. Although the average rate was lower under FI filtered light, this difference was not statistically significant due to the large variation within the treatment. **C**) Rate of transpiration under the FI filter relative to the unfiltered control. The FI treatment was significantly lower than control (p < 0.01). **D**) Photo of representative tomato plants at 30 days.

### Anthocyanin content was significantly higher under the FI filter in lettuce

Certain secondary metabolites with photosynthetic relevance were extracted from lettuce leaf tissue to look for acclimation to the altered light environment under the OSC filters. Chlorophyll, for example, has been shown to increase in overall concentration in response to a lower R/B ratio (34). There were no significant differences in the ratio of chlorophyll *a* to *b* (Figure 3). Surprisingly, this was also the case when light intensity as well as spectra was varied (32), despite a well-documented correlation between lower light intensity and an increase in chlorophyll *b* (35, 36). However, anthocyanins that protect the plant by absorbing excess light varied between treatments. The FI treatment had a significantly higher anthocyanin content than all other treatments (Figure 3). Despite receiving approximately the same amount of light, the lettuce plants in this treatment accumulated more anthocyanins than those under the other filters or the white light control. While anthocyanin content in lettuce has been shown to increase with exposure to UV light (27), increased blue light (37) and increased light intensity (38), the differences in blue light and total light intensity between the FI treatment and the other treatments were small (∼15 μmol m^-2^ s^-1^ blue and ∼70 μmol m^-2^ s^-1^ TPFD), and UV light was negligible in all treatments. This suggests that aspects of the FI transmission spectrum enhance anthocyanin content beyond what could be expected from the amount of blue light it transmits.

### Gene expression networks identified genomic responses

A transcriptome analysis was conducted for each species to detect gene expression differences caused by the variation in filter spectra and light intensity in lettuce and tomato leaf tissue. Such analyses are often used to identify emergent traits caused by the combination of changes in the expression of the tens of thousands of genes in the genome that are not immediately apparent in the phenotype. To examine global patterns of gene expression in response to the filters, we used a network-based approach to identify distinct groups of co-expressed genes that shared a similar relationship with the filter-specific light spectra and were consistent when light intensity was varied. Gene networks were individually constructed for each species using an unsupervised machine learning approach. Datasets collected under varied TPFD were included to differentiate between genomic responses to altered spectrum from those caused by light intensity changes. The resulting networks were used to build a common consensus network for each species containing clusters of genes (i.e. modules) that are shared between the intensity-independent and intensity-dependent networks. The consensus networks identified 35 modules ranging in size from 39 to 3,656 genes in the cluster dendrogram for lettuce (Figure S6) and 43 modules ranging from 46 to 2,326 in tomato (Figure S7).

Genes within a given module are considered to have highly similar expression profiles. Each module is described by a single value called the module eigengene that represents the expression profile of the entire module. These module eigengenes were used to assess relationships between gene expression profiles and quantitative measures of light quality, such as TPFD, red to blue (R/B) ratio, red to far-red (R/FR) ratio. We identified clusters of genes that responded similarly to these parameters by comparing the sign of the correlation between the module eigengene and each quantitative measurement. These clusters or modules are unique to each species.

In lettuce, Module L2 (2,397 genes) showed a strong positive correlation with TPFD (*r*=0.9, p=1e-07, Figure 5), indicating that genes in this module are strongly influenced by small increases in light intensity. Module L12 (3,656 genes) also showed a strong correlation with TPFD but in the opposite direction (*r*=-0.8, p=3e-05). This suggests an inverse relationship between genes in Module L2 and Module L12, where higher values of TPFD are associated with higher expression of genes in Module L2 but lower expression in Module L12. The same two modules also had opposite correlations with the R/FR ratio. Module L2 was negatively correlated with the R/FR ratio, meaning that these genes tended to have higher expression when lettuce was grown under a lower R/FR ratio (*r*=-0.9, p=3e-06). However, when light intensity varied in addition to the spectrum, the correlation between Module L2 eigengene and the R/FR ratio was not significant (p=0.05). A similar pattern was seen in Module L12 where there was a positive correlation with the R/FR ratio (*r*=0.8, p=1e-05).

**Figure 5.**
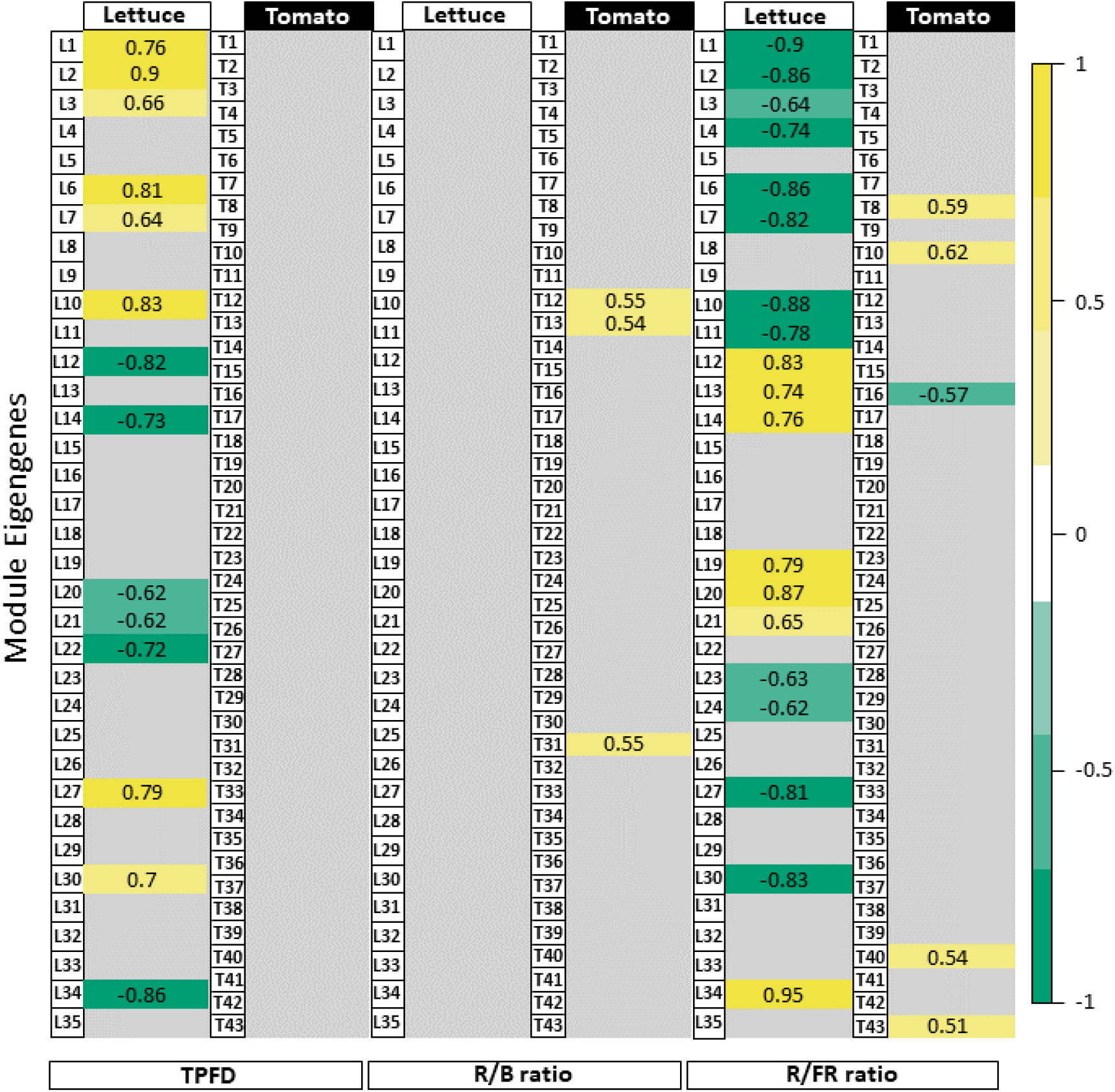
Module-trait relationships with selected light traits. Table of correlations between measured light parameters and the 35 lettuce modules and 43 tomato modules of genes identified in the network analysis. Strong positive correlations are dark yellow while strong negative correlations are dark green. Values given are the correlation coefficient (*r*) for each relationship. Although gene expression in lettuce varied in response to changes in TPFD, the same responses were not seen in tomato. The responses to light quality traits (i.e., R/B and R/FR ratios) were similarly variable between lettuce and tomato. Tomato gene modules correlated more strongly with the R/B ratio while the R/FR ratio had more influence on gene expression in lettuce. Modules showing weak or no significant correlation with a trait were filtered by applying a p-value threshold (p<1e-05). See Figures S8,11-13 for the complete module-trait relationships for lettuce and tomato.

In tomato, 43 modules of genes with similar expression were identified (Figure 5). Surprisingly, the correlations between several modules and TPFD observed in lettuce were not observed in tomato. Rather, there were no statistically significant correlations between any modules of genes and TPFD when tomato plants were grown under OSC filter with the same light intensity. This suggests that the small differences in TPFD were not sufficient to modify gene expression in tomato. The lack of correlation with TPFD indicates that the experimental design of the tomato experiment was optimized by reducing the already small light intensity variation in the lettuce experiment. The tomato treatments are closer to each other in TPFD, which allowed for a clearer interpretation of the filters’ spectral effects on gene expression. Although gene expression of very few modules correlates significantly with the spectral traits, the correlations that do exist offer insight into the plant’s response to the modified OSC spectra. In particular, the representative eigengene of Module T12 (219 genes) had a positive correlation with the ratio of R/B light (*r*=0.55, p=0.02). This indicates that an increase in R/B ratio, such as that produced by the FP filter, increases expression of the genes assigned to this module.

### Hub genes in lettuce and tomato

We focused on specific genes with expression patterns that were highly correlated with a particular module and its representative eigengene. In lettuce, Modules L2 and L12 were selected to investigate the R/FR dependence. One such hub gene in Module L2 is *elongated hypocotyl 5* (*HY5)*, which encodes a light-responsive transcription factor important for photomorphogenesis (23). As expected of a hub gene, *HY5-1* expression closely matched the pattern of both Module L2 eigengene expression and the overall expression of all Module L2 genes (Figure 6). Accordingly, *HY5* expression was also negatively correlated with R/FR, suggesting that *HY5* expression increases in response to lower R/FR ratios. Indeed, the treatment with the lowest R/FR ratio (Data S1), also had the highest expression of *HY5-1*. Conversely, a *high affinity nitrate transporter 2*.*1* gene (*NRT2*.*1-1*, LOC111883156), a hub gene of Module L12 (kME=0.9, p=2e-06), had an opposite expression pattern with the lowest expression in FI, compared to the other treatments. Surprisingly, HY5 has been identified as an enhancer of NRT2.1 in roots and is expected to increase, not decrease, expression of its target, a gene important for nitrate uptake and nutrient acquisition (24).

**Figure 6.**
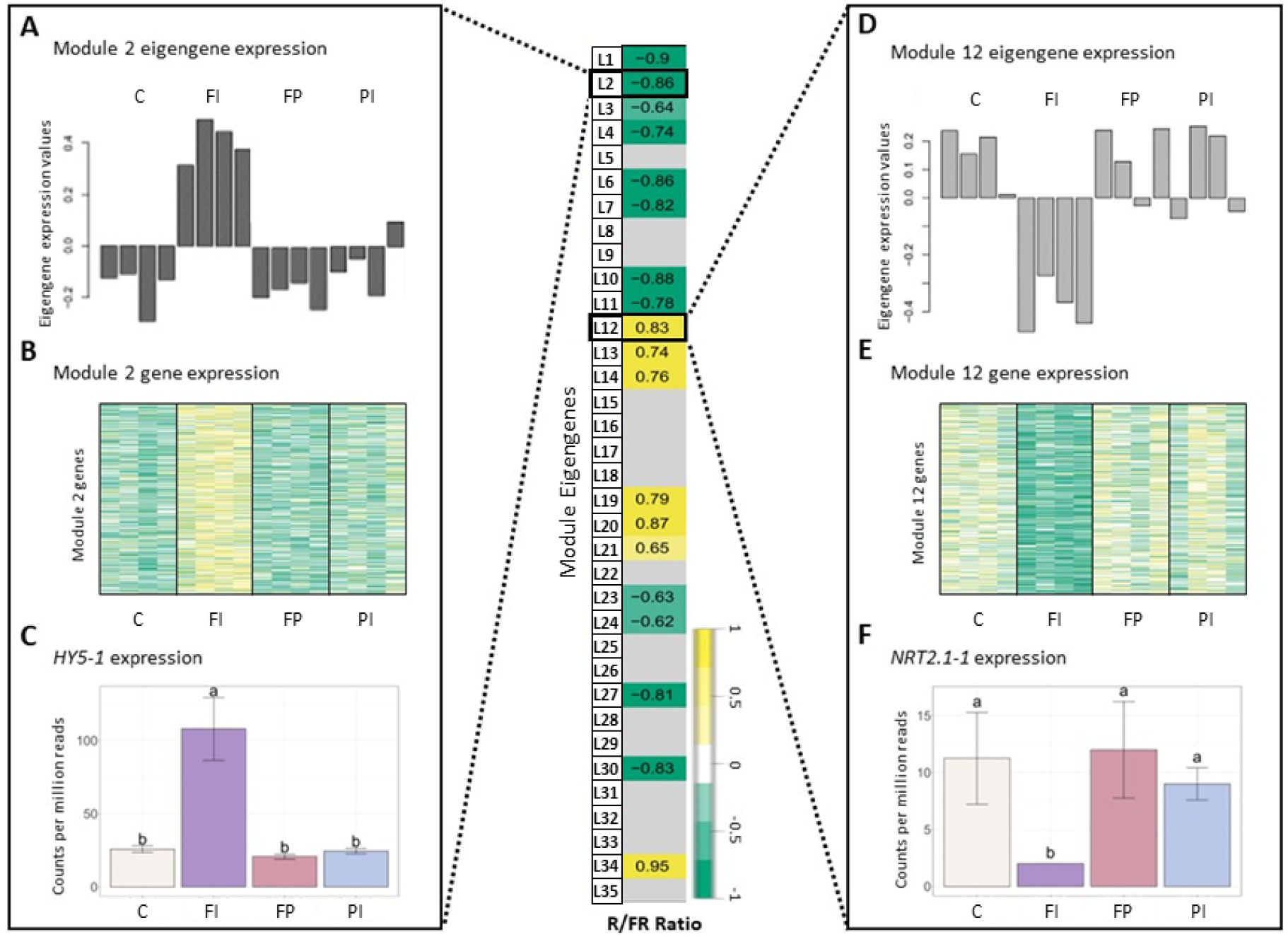
Gene expression in lettuce correlated with R/FR ratio. Module-trait relationships and differential gene expression between lettuce filter treatments and control. **A**) Normalized expression of the Lettuce Module 2 eigengene expression for each of the treatments by sample. Table values given are the correlation coefficient (*r*) for each relationship. Insignificant correlations were removed (p<1e-05). Error bars indicate the standard deviation. Letters indicate significance from ANOVA and Tukey test (p<0.05). See Figure S11 for full table. **B**) Heatmap of gene expression for all genes in Lettuce Module 2, where dark yellow indicates a strong increase in expression and dark green indicates a strong decrease in expression. **C**) Normalized gene expression of a Lettuce Module 2 hub gene, *elongated hypocotyl 5* (*HY5-1*). **D**-**F**) Eigengene expression, gene expression and hub gene expression (*high-affinity nitrate transporter 2*.*1, NRT2*.*1-1*) for Lettuce Module 12. The treatment with the lowest R/FR ratio, FI, had higher gene expression in Lettuce Module 2, resulting in a negative correlation with the R/FR ratio. In contrast, Lettuce Module 12 had a positive correlation with R/FR ratio due to the downregulation of expression in FI, relative to the other treatments.

A gene that negatively regulates flowering in tomato, *self pruning 5G* (*SP5G*) (19), had a strong correlation with the Module T12 eigengene (kME=0.9, p=3e-07) and was, therefore, a good candidate hub gene (Figure 7). This module had a positive correlation with the R/B ratio. Expression of *SP5G* increased under the FP filter, which had a relatively high R/B ratio (4.5), and decreased under the FI filter with its relatively low R/B ratio (2.8), although this decrease was not statistically significant. The same correlation between Module T12 and the R/B ratio was not seen when light intensity varied (Figure S8). While FP had the highest expression levels as expected, *SP5G* expression in the FI treatment was not significantly lower than the high light control.

**Figure 7.**
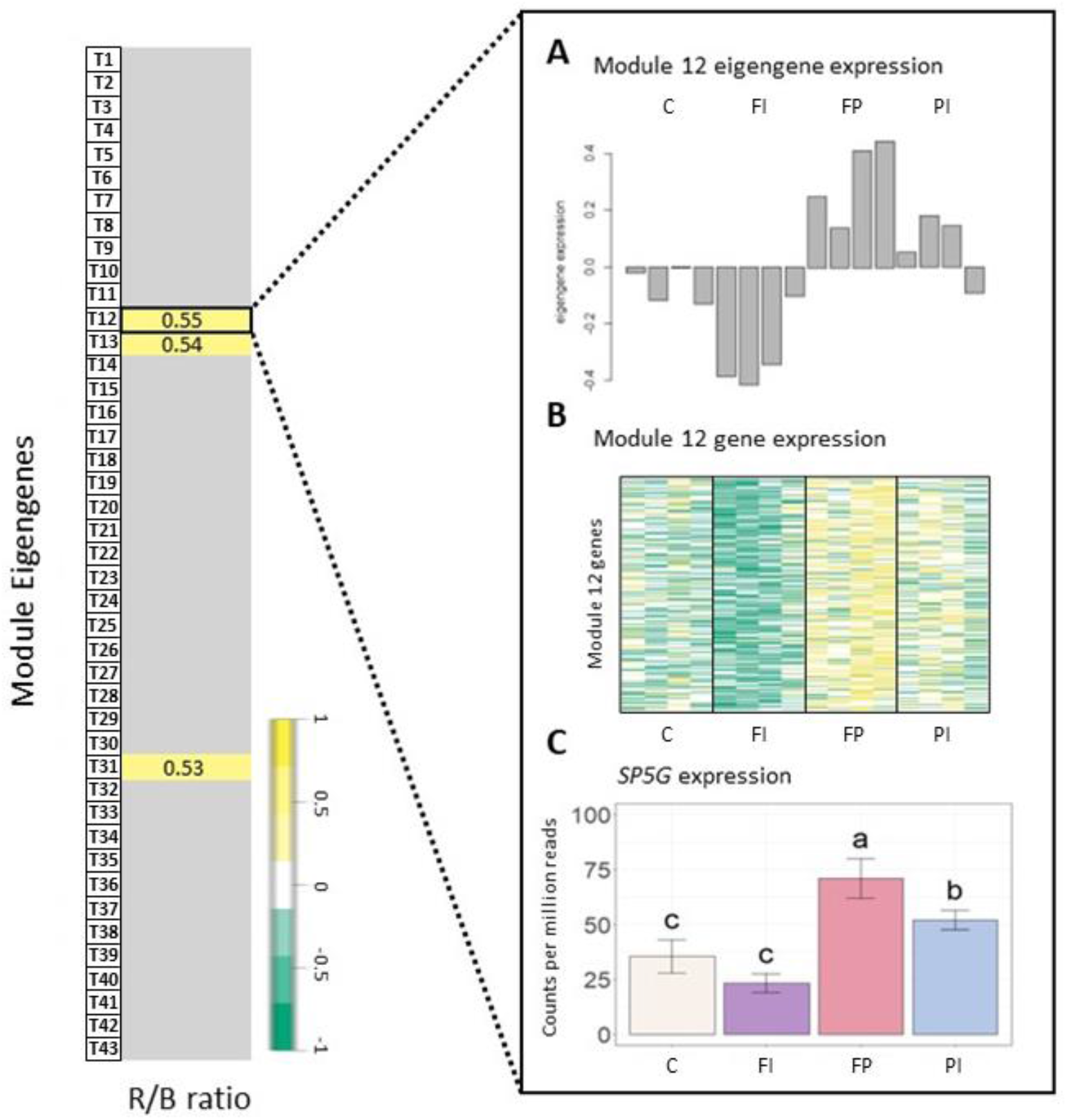
Gene expression in tomato correlated with R/B ratio. Module-trait relationships and differential gene expression between tomato filter treatments and control. **A**) Normalized expression of the Tomato Module 12 eigengene expression for each treatment by sample. The correlation coefficient (*r*) is given for each relationship. Insignificant correlations were removed (p<0.05). See Figure S12 for full module-trait relationships table. **B**) Heatmap of gene expression for all genes in Tomato Module 12, where dark green indicates a strong increase in expression and dark yellow indicates a strong decrease in expression. **C**) Normalized gene expression of a Tomato Module 12 hub gene, *self pruning 5G* (*SP5G*). The treatment with the highest R/B ratio, FP (FTAZ:PCBM filter), had higher gene expression in Tomato Module 12, resulting in a positive correlation with the R/B ratio. This correlation is exemplified in a hub gene of Tomato Module 12: *SPG5*. Error bars indicate the standard deviation. Letters indicate significance from ANOVA and Tukey test (p<0.05).

The pattern of expression of *SP5G* corresponds with the timing of flowering observed. The FI treatment had lower expression of this floral repressor (Data S3). Although flowering data collection was limited by the size restrictions within the growth boxes, our observations indicate that the FI treatment flowered earlier than the control treatment (Data S2). In contrast, the FP treatment had high expression of *SP5G* and indications of delayed flowering, relative to the control (Data S2, S3). *SP5G* is thought to be downstream of both the red light-sensitive photoreceptor, phyB1 (39), and the blue-light sensitive photoreceptors, cry1a and cry2 (40). This provides a potential mechanism for the sensitivity of this gene to the differing R/B ratios in the filter treatments. The characterized relationship between phyB1 and *SP5G* expression also suggests a correlation between gene expression and the R/FR ratio that directly impacts the activation state of phytochromes. A lower R/FR has been shown to increase expression of *SP5G* and delay tomato flowering (39). While the network analysis did identify a correlation between *SP5G* expression and R/FR ratio, it was a weaker correlation with the opposite sign expected from the literature (GS=0.58, p=0.02).

### Gene expression patterns vary between lettuce and tomato

A differential gene expression analysis was performed to contrast each filter treatment against its respective control to quantify changes in the expression of individual genes. In lettuce, few differences were observed for all filter treatments relative to the control except FI. There were 1,143 DEGs identified in the FI/C comparison but only 15 DEGs in FP/C and 12 DEGs in PI/C (Figure 8a). While filter degradation did result in changes in TPFD by the end of the experiment (Figure S3), this small variation in light intensity alone can hardly account for the large differences between the FI treatment and the other treatments. As reported earlier, gene expression of several large modules of genes correlated with the R/FR ratio as well as TPFD in this experiment. This suggests that the R/FR ratio and possibly other spectral characteristics unique to the FI filter resulted in differential gene expression. In particular, *HY5-1* gene expression was increased by a log2-fold change of 2.2 in FI, which is consistent with a previous study that found that *HY5* expression increased with lower R/FR ratios (41). Because HY5 is a transcription factor that controls expression of many downstream genes, it is not surprising that many of these genes were also differentially expressed in the FI treatment. Among these were several anthocyanin modification enzymes that were upregulated relative to the control (Table S1). The increase in expression of anthocyanin modification genes correlates with the increase in anthocyanin concentration seen in Figure 3, where the FI treatment had a significantly higher concentration than all other treatments. Other genes known to be directly or indirectly regulated by HY5 were also upregulated, including several related to nitrate uptake, defense against predation and the circadian clock.

**Figure 8.**
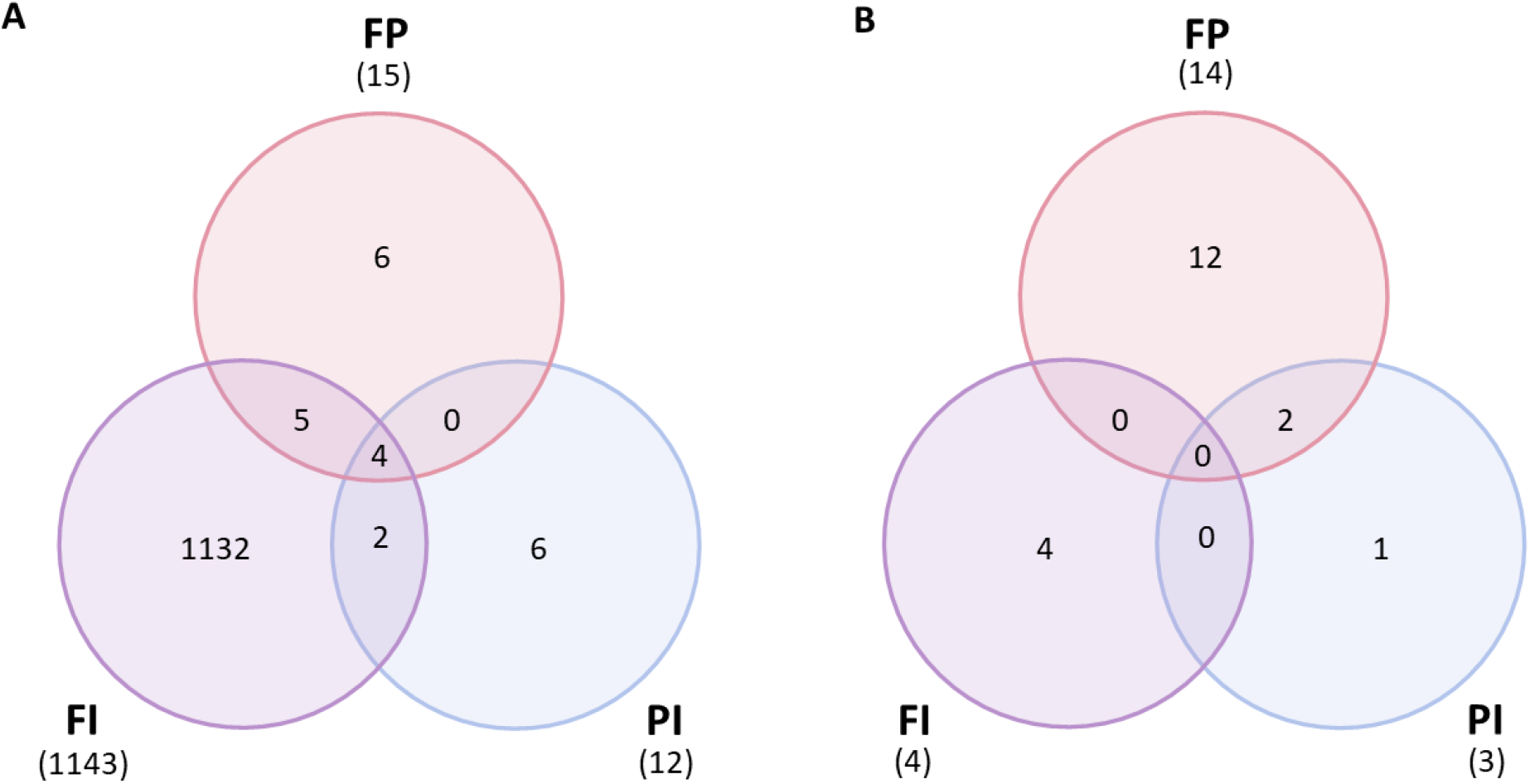
Differentially expressed genes between filter treatments and control. **A**) Lettuce normalized gene expression (counts per million) contrasted between each treatment for each filter (FI, FP and PI) and the control (C) (p<1e-05). Lettuce grown under the FI filter had thousands of differentially expressed genes relative to the control, while the FP and PI treatments both had fewer than twenty DEGs each. **B**) Tomato normalized gene expression (counts per million) contrasted between each treatment for each filter (FI, FP and PI) and the control (C) (p<1e-05). Very few genes were differentially expressed between each light treatment in tomato, with the majority found in the FP/C contrast.

However, there were very few differences between the tomato filter treatments and the control. There were only 4 DEGs in the FI/C contrast, 3 in the PI/C contrast and 14 in the FP contrast (Figure 8b). Unlike in lettuce, the FI filter did not cause a dramatic expression profile difference relative to the control. The FP filter had a greater impact on the tomato genome, although this was only a few genes more than either FI or PI.

Many of the differentially expressed genes observed in the lettuce experiments were not seen in tomato. The expression of the transcription factor, elongated hypocotyl 5 (HY5-1), was upregulated in lettuce plants grown under the FI filter. In particular, the FI treatment had higher *HY5-1* expression than all other treatments. HY5 mediates light-responsive signals by regulating a diverse range of downstream genes, including *high-affinity nitrate transporter 2*.*1* (*NRT2*.*1-1*) (42). The interaction between light, HY5 and downstream targets such as *NRT2*.*1-1* points to modification in nutrient uptake and accumulation driven by the spectrum FI OSC filter in lettuce. However, this interaction was not observed in tomato. Expression of *HY5-1* was not significantly higher in the tomato FI treatment compared to the control (Figure S9). Furthermore, there were zero counts of *NRT2*.*1* expressed in sampled tomato leaf tissue. It seems likely that the expression of this nitrate transporter is restricted to root tissue in tomato, which was not analyzed here.

Among the genes that were differentially expressed between OSC filter treatments and the white light control in tomato, the flowering regulators *self pruning 5G* (*SP5G*) and *GIGANTEA* (*GI*) were upregulated in FP and both FP and PI, respectively (Figure S9; Table S2). GI, in particular, plays many roles in regulating the circadian clock and various light-responsive processes and is downstream of phyB (43). The expression of *GI* increased with both the high relative amounts of blue light in the PI filter and the high relative amounts of red light in the FP filter. Additionally, genes involved in diverse pathways such as *response to low sulfur 3-like, inositol-1,4,5-triphosphate-5-phosphatase* (*5PT1*) and *terpene synthase* group gene, *TPS12*, were differentially expressed in response to spectral changes alone (Figure S9).

The expression of the photoreceptors and downstream genes in tomato had patterns different from those seen in lettuce as well (Figure 9). While the activity of photoreceptors as enzymes post-translation is most commonly discussed in studies on plant light responses, photoreceptor gene expression is also responsive to light conditions (44-47). The gene expression of *phytochrome A* (*phyA*), which encodes one of the red light photoreceptors, had the most variation in lettuce and tomato. Phytochrome expression is known to increase in response to high levels of red light, improving the plant’s ability to perceive these wavelengths of light (46). Accordingly, this gene was more highly expressed in the FP treatment, although only in tomato. There was less variation in the gene expression of other photoreceptors, which also have a wavelength-sensitive increase in expression (44, 47).

**Figure 9.**
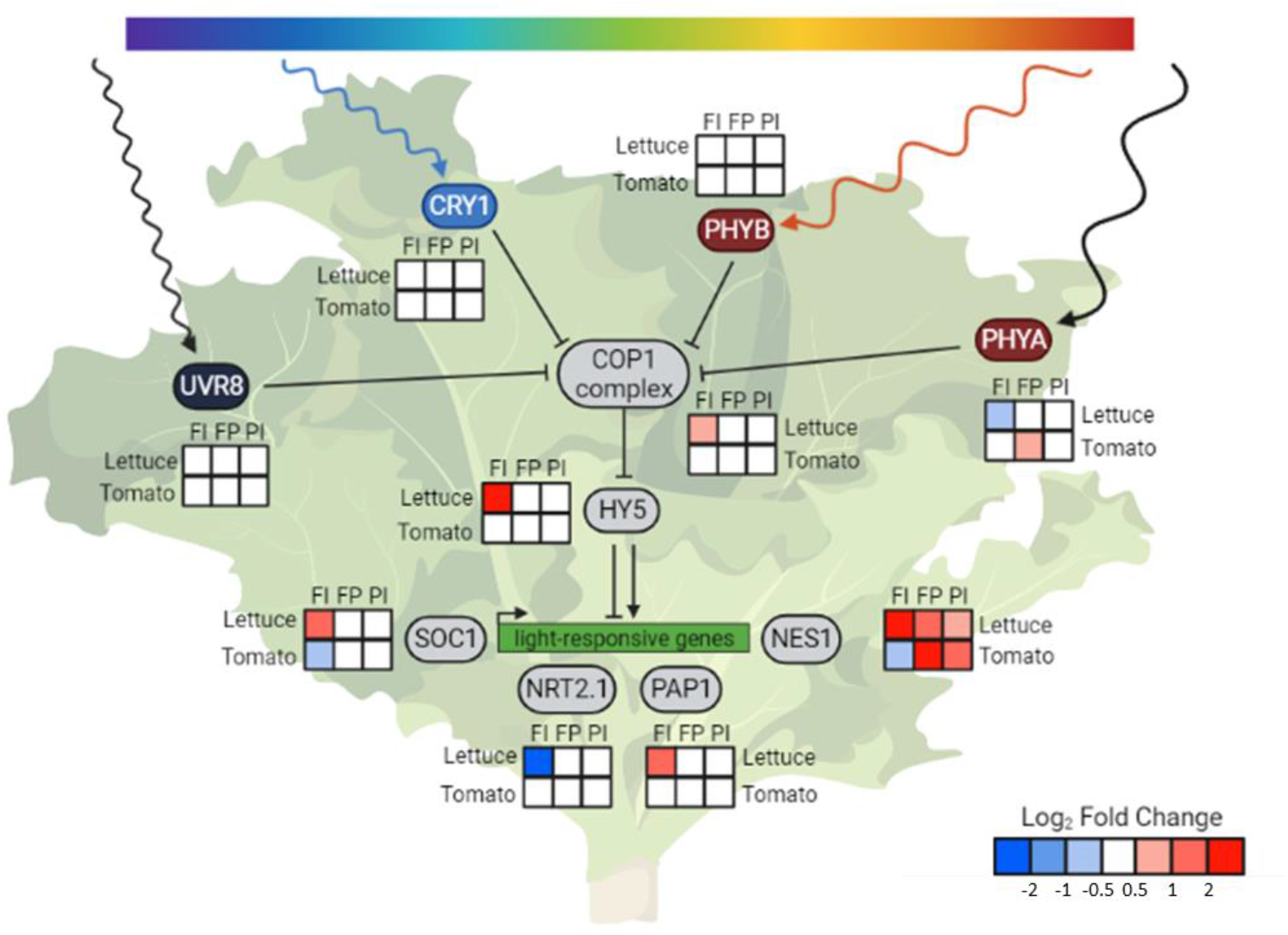
HY5 integrates light signals and regulates downstream genes. Condensed photoreceptor signaling pathway with expression of downstream genes of interest in lettuce and tomato. The photoreceptors indirectly upregulate expression of *HY5* through inhibition of the constitutive photomorphogenic 1 (COP1) complex formation. Elongated hypocotyl 5 (HY5-1) is a transcription factor that is a known or hypothesized regulator of many light-responsive genes, including *suppressor of overexpression of CO 1* (*SOC1*), *high affinity nitrate transporter 2*.*1* (*NRT2*.*1-1*), *production of anthocyanin pigment1/MYB75* (*PAP1*) and *(3S,6E)-nerolidol synthase 1* (*NES1-1*). *COP1* itself was differentially expressed in response to light quality changes in lettuce but not tomato. The changes in *COP1* expression are in the opposite direction of many of the trends observed in *phyA1* expression, indicating gene-level regulation of this gene. Despite a lack of differential expression of *COP1* and *HY5*, expression of downstream genes, such as *NES1*, were altered in tomato, indicating another method of light regulation. Relative transcript abundance is represented by log_2_ fold changes between specified treatments. Created with BioRender.com.

The phytochromes, along with the cryptochromes, inhibit the formation of the complex that COP1 ubiquitin ligase forms with suppressor of phyA 105-1 (SPA1), which prevents the degradation of transcription factors such as HY5 and leads to photomorphogenesis (23). Our analysis revealed that, in addition to the photoreceptors, *COP1* itself was differentially expressed in lettuce in response to both light quality changes and intensity changes. The changes in *COP1* expression are in the opposite direction of many of the trends observed in *phyA* expression. This may indicate that the expression of *COP1* in lettuce, in addition to activity of its protein, is regulated by light. The expression of several genes downstream of COP1 was also altered in response to changes in spectrum and the overall amount of light in lettuce. One of these genes is *suppressor of overexpression of CO 1* (*SOC1*), a transcription factor that regulates flowering and has been identified as a putative target of HY5 (48, 49). Production of anthocyanin pigment 1/MYB75 (PAP1) is a transcription factor that regulates anthocyanin accumulation, and whose gene expression is also directly regulated by HY5 (42). Accordingly, *PAP1* expression mirrored the trends seen in *HY5*. This provides a genomic basis for the increase in anthocyanin content seen in the lettuce FI treatment.

While many of these downstream changes were seen only in lettuce, altered expression of a volatile-producing gene was observed in both species (Figure 7). *(3S,6E)-nerolidol synthase 1* (*NES1-1*) gene expression increased in response to both light intensity and altered spectrum. While it has not been demonstrated that *NES1* is directly regulated by HY5, it is a known regulator in the terpenoid synthase pathway of which *NES1* is a member (50). *NES1* encodes an enzyme that produces nerolidol, a volatile compound released by the plant in response to wounding by herbivorous insects. This molecule has been shown to attract predator insects in tea plants, protecting the plants from further damage by reducing the number of pests (51). In lettuce, *NES1* was upregulated in response to all three OSC filters. In tomato, only the FP and PI filters resulted in an increase in *NES1* expression, while the FI treatment showed lower expression.

## Discussion

Our experiments were designed to account for both the quantitative and qualitative aspects of OSC-filtered light on lettuce and tomato as representative crops. Despite the morphological similarities between treatments, the physiology of the plants was altered by the differences in light quality under the ST-OSC filters on a molecular and transcriptomic level. These molecular tools enable us to predict physiological responses that can now be tested to further improve the productivity and sustainability of crops to be grown in ST-OSC greenhouses and ultimately breeding or engineering of crops to specifically optimize their performance.

### Transcriptome analysis identified physiological and metabolic changes

The transcriptome analysis was a screening tool for the many aspects of plant growth and development that could not be measured directly. Beginning with the photoreceptors that sense light, expression of many genes was altered by changes in either light intensity or light quality, or both (Figure 9). The expression patterns seen in the photoreceptors reflect the light environment of the plants. There was minimal variation in *UVR8-1* expression and negligible UV light, while there was greater variability in *phyA* that corresponded to the variability in red light between treatments (Table 1). A common theme of the transcriptome analysis in lettuce was the differential expression of *HY5*, a light-regulated gene that encodes a transcription factor that controls the expression of many genes that drive photomorphogenesis and is regulated by the COP1 complex. *HY5* was differentially expressed in response to altered spectrum and altered light quantity only in lettuce, leading to many downstream changes. Tomato lacked this differential expression of *HY5*.

### Impact of OSC-altered spectra on flowering

The upregulation of *SOC1* in lettuce grown under the FI filter relative to the control with similar light intensity suggests that this spectrum may trigger an earlier flowering time. The regulation of flowering is an important aspect of crop development that can impact harvest. In particular, the prevention of early flowering in lettuce can improve harvests by extending the growing period before the lettuce begins to bolt, altering its metabolite and flavor profile (52). Although this would be undesirable in several crop species, this effect was not seen in the other OSC filters tested, which yielded similar biomass. This is an example of the importance of considering the transmission in the 400-750 nm range when selecting OSCs for greenhouses.

In tomato, probable transcription factor SP5G is a known repressor of the start of flowering (39). Despite timing of the RNA tissue sampling for the genomic analysis after the initiation of flowering, elevated expression levels of *SP5G* were detected in combination with the delayed flowering in the FP treatment. The light spectrum of the FP filter is similar to the spectrum of the commercial OSCs used in a greenhouse tomato experiment (32). The tomato plants in this experiment also experienced a reduced amount of blue light relative to red, in addition to a decrease in the R/FR ratio. Although the authors did not report a delay in flowering, the first three fruit harvests of the indeterminate tomato plants were lower in yield compared to the control. Taken together, this suggests that the FP filter and other OSC filters with similar spectral profiles may have a negative effect on flowering crop production, similar to the potential negative impact on bolting in lettuce associated with the FI filter. On the other hand, FI, the treatment with a more balanced B/R ratio that flowered one day before the control had the lowest levels of *SP5G*. Although *SP5G* was not significantly differentially expressed between this treatment and the control, this may indicate that the FI filter still holds some advantage over others in promoting beneficial agronomic traits in tomato and similar high-light species.

### Impacts from OSC spectra on lettuce nutritional content

Crops rich in anthocyanins are more nutritious and have a positive effect on human health when included in the diet (53). Anthocyanin content in red cultivars of lettuce has been shown to increase in response to increased light intensity (38) and altered spectrum (37, 54). The increase in *PAP1* expression provided a genomic and molecular basis for the increase in anthocyanin content seen in the lettuce FI. While many of the genes in the anthocyanin biosynthesis pathway were not differentially regulated on a statistically significant scale, several genes involved in the modification of anthocyanins into more stable forms were upregulated (55) (Table S1, Data S3). This suggests that the primary mode of anthocyanin accumulation in lettuce may have been through the stabilization of anthocyanins rather than novel biosynthesis. Although the lettuce FI treatment did have a somewhat higher amount of light than other PC treatments, there is reason to believe that the increase in anthocyanins could be the result of the FI spectrum alone. The FI treatment had a wavelength profile with higher amounts of blue and red light and lower amounts of green relative to the other treatments (Table 1), and anthocyanin content has been shown to increase under red and blue light (37).

While nitrate is required for growth and must be supplied in fertilizer, high nitrate content in lettuce leaves is an important consumer concern with human health effects (56). Several nitrogen transporter genes that allow the plant to take up nitrogen from the soil and move it within the plant were differentially expressed in the FI treatment relative to control, including *NRT2*.*1-1* (Figure 9; Table S1). Surprisingly, expression of *NRT2*.*1* was downregulated when *HY5* was upregulated, although HY5 is typically considered to be a positive regulator of *NRT2*.*1* (24). A recent study has demonstrated that HY5 can act as a negative regulator of *NRT2*.*1* under certain conditions (41). It should also be noted that much of the research on NRT2.1 is focused on nitrate uptake from the soil and is primarily analyzing expression in root, not leaf, tissue. The downregulation of nitrate transporters in leaf tissue may indicate that nitrate is selectively reduced in the roots instead of the leaves and accumulates less in the leaves. Additionally, the decrease in *nitrate reductase* (*NR*) expression could indicate lessened demand for the conversion of nitrate (NO-3) to nitrite (NO-2) in the leaves (Figure S10). These changes may correlate with desirable low nitrate levels in leaf tissue and improved nutrition. Nitrate levels in lettuce have been shown to vary in response to different light intensities (57). Nitrogen use efficiency and fertilizer requirements may also be affected. Because fertilizer uptake and nitrogen content were not quantified in this study, further experimentation is needed to confirm these changes.

A limitation of our experimental design was the variation in light intensity between treatments, especially in the lettuce experiment. Although this is not unlike real world variation that would result from differences in filter transmission or varied shading from bench to bench in the greenhouse, the changes in TPFD made it more difficult to distinguish between changes that were solely due to the filter spectra. A combinatorial study, where treatments are compared to others that vary in only filter or TPFD and not both at once, would more clearly distinguish between light intensity and light quality effects. Such a study could better predict the changes in crop growth and yield caused by general OSC shading as well as the effects that would vary based on the transmission of the active layer. The many potential metabolic changes that were suggested by differential gene expression but not quantified, especially changes in nitrogen acquisition, fruiting and herbivory defense in other important greenhouse crops should be investigated. Additionally, recent modeling of plant growth and energy harvesting in OSC-greenhouses have identified a number of promising organic semiconductor active layers, which includes the FTAZ:IT-M (FI) and PTB7-Th:IEICO-4F (PI) systems studied here (30). Further study of OSC active layers, particularly under natural sunlight, would validate the model’s ability to identify materials that can produce economically viable crops. More broadly, a combinatorial transcriptomic approach to the study of plant light responses in general could yield valuable insight into the signal integration between different aspects of light quality and light intensity.

### Future directions

Our experiments presented here offer molecular insight into plant growth and development under OSC filters. We found no negative impacts on the accumulation of biomass or on the quantified secondary metabolites when light intensity was controlled. The differential gene expression, especially the upregulation of key regulator genes under the FI filter in lettuce and the FP filter in tomato, is worthy of further study to discover how these changes translate to important aspects of crop production. In particular, the gene expression changes related to the initiation of flowering pointed to economic impacts for crops like lettuce, where early flowering can damage harvests by introducing a bitter taste, and tomato, where improved fruit development can increase yields. The advantage of a transcriptome analysis in the study of OSC-grown plants is that these key modifications can be identified without the need to directly measure each aspect of plant growth and development.

A major limitation to commercialization of ST-OSCs in greenhouses is filter fading. This issue is currently being addressed, and we expect to develop commercially meaningful lifetimes for OSC filters in the near future. This will enable scale-up production and commercialization. To have a meaningful impact on sustainable food production, these ST-OSC greenhouses also need to be able to produce a larger variety of crops in different climate zones. While we have modeled the potential for economic value with some crops (30), lower material costs and higher efficiency will provide a path to not only grow locally desirable vegetables from strawberries and beans to eggplants, but also row crops such as corn, wheat and root vegetables. Most of the improvements are expected to come from OSC efficiency increases, better materials and different systems like flexible OSC shades that can be used in greenhouses in a similar, more temporary way as current shading techniques and paint are applied.

In addition to the material science and battery storage improvements that can be anticipated, breeding or genetic engineering of crops that are specifically adjusted to these modified growth conditions can be considered. Many of the traits that evolved through natural selection or were bred for field crops are no longer required in controlled environment agriculture. Stress response mechanisms in plants for survival in varying biotic and abiotic environments often reduce yield. When the environment can be controlled, those yield-reducing stress responses are no longer required and can be removed through conventional breeding or engineering (58).

The magnitude of intensification of agricultural crop production in these net zero energy greenhouses would not only contribute to an increased food demand even on marginal land, but it can also spare land, so it can be converted to other ecosystems, which could provide income from carbon credits through reforestation or other ecosystem services. Future research focuses on improving ST-OSC stability and testing of other crop systems for productivity and sustainability traits in ST-OSC GreenERhouses.

## Materials and Methods

### Experimental Design

Red oak leaf lettuce (*Lactuca sativa*) was grown with adjustments made to the height of the growth boxes so that each treatment received similar light intensity in addition to the consistent height, variable intensity setup previously described (33). Eight lettuce plants from each treatment were harvested at 21 days post germination (transplant stage) and the remaining eight plants from each treatment at 35 days post germination (harvest stage). Four plants from each harvest per box were used for biomass measurements, while the remaining four were used for tissue sampling. Tomato plants (*Solanum lycopersicum* cv. Moneymaker) were grown under these conditions with modifications. Seeds were sown on rockwool and germinated in the same growth chamber with metal halide and incandescent lighting to approximate natural sunlight. Eight seedlings of uniform size and age were selected and transplanted into individual blocks of larger rockwool and moved inside the treatment boxes with nearly 100% OSC roof coverage. Tomato plants were harvested after flowering, 30 days past the two-leaf stage when plants were moved under the filters.

The growth boxes were covered with either an OSC filter, a clear glass or a shaded control on top to simulate a greenhouse roof. The positions of the rockwool blocks were rotated to avoid positional light effects as in the lettuce experiment. The consistent light intensity between treatments allowed for comparison of the influence of the light spectra on plant physiology. The results of these experiments were compared to the previously reported experimental design where all filters were positioned at a consistent height to model the roof of a greenhouse and therefore produce different TPFD due to the differences in filter transmission. Five replications of the lettuce experiment with consistent light intensity were conducted. One replication was conducted of both tomato experiments, using consistent and variable light intensity. Lettuce light conditions were measured as previously reported. Reported percent colors for tomato were measured using a spectrophotometer (Black Comet-SR, Stellar Net, Inc., USA) except the high light control treatment, which was assumed to have the percent colors of the low light control. PPFD was measured using a quantum sensor (LI-190R, LI-COR, Inc., USA) and TPFD was calculated from PPFD and percent colors.

### Filter fabrication

OSC filters were made as previously described (33). Solutions of the organic semiconductor active layers were wire bar-coated onto glass substrates. A second sheet of glass was adhered under heat to each of the glass substrates with ethylene vinyl acetate films for encapsulation. Optical epoxy (Norland 63) was cured around the edge of the filter stack as an additional seal. Twelve of these filters, each 20×10cm, were arranged in a single layer above a layer of PEDOT:PSS (PH1000, Hareus) coated onto a PET substrate to simulate the transmission of full OSC devices for each filter treatment.

### Biomass measurements

Measurements of fresh weight, dry weight, leaf area and leaf number were collected as previously described at Day 21 and Day 35 after germination for lettuce and at Day 30 for tomato (33). The 21-day early harvest corresponds to the age when young lettuce is typically transplanted. Both fresh and dry weights are above ground measurements that do not include root tissue. Dry weight was measured after leaves were dried at 65°C for three days. Leaf area was measured by leaf meter (LI-3000, LI-COR, Inc., USA) and summed per plant. Leaf number was also summed by plant and excluded any emerging leaves less than 1 cm in length. Additional measurements were taken of the tomato plants. Height was measured from the top of the rockwool block to the highest point of the plant. Visible flower buds and open flower buds were recorded per plant after initiation of flowering and at harvest. The number of leaflets was counted per plant. Height and leaf number were collected for all eight plants in each treatment. All other biomass measurements were collected for the four tomato plants per treatment not used for tissue sampling.

### Extraction and quantification of secondary metabolites

Secondary metabolites were extracted from ground frozen lettuce leaf tissue as previously described (33, 59). Leaf tissue was collected from four harvest stage plants and immediately frozen in liquid nitrogen and stored until ground. Ground frozen leaf tissue was weighed and suspended in extraction buffer. A BioTek Synergy HT microplate reader (BioTek Instruments, USA) was used to measure absorbance.

### Photosynthetic data collection

A LI-6400XT (LI-COR, Inc., USA) was used to collect photosynthetic data from lettuce as previously described (33). Two sample measurements were collected per leaf, two leaves per plant and four plants per treatment beginning five days before the final harvest. Photosynthesis was measured *in situ* inside the growth boxes to observe the impact of the light intensity and spectrum created by the OSC filters. The chamber door was kept closed during data collection to minimize changes to the environment and a black cloth was used to block ambient white light from entering around the equipment. A CO2 scrubber was used to prevent elevated CO2 levels from researcher exhalation. PPFD was monitored as measured by the instrument to ensure lighting conditions remained consistent throughout the data collection for each treatment. Photosynthetic data collection was limited in tomato due to experimental constraints.

### RNA extraction

Mature leaf tissue was collected from four plants per treatment in one replicate and ground in liquid nitrogen. The PureLink RNA Mini on-column kit with TRIzol (ThermoFisher Scientific, Inc., USA) was used to extract total RNA. An on-column DNAse treatment with additional off-column DNAse I treatments were used to remove DNA contamination. The mRNA library preparation and sequencing were performed by BGI Genomics Co., Ltd. (Shenzhen, China) with polyA selection by an oligo dT library. All 32 samples were multiplexed, pooled and loaded together. Sequencing was conducted on a DNBSEQ™ Technology Platform.

### Transcriptome analysis

Raw reads were checked for quality standards using FastQC (v. 0.11.9) (http://www.bioinformatics.babraham.ac.uk/projects/fastqc/) and only high-quality read pairs (base score above Q30) were subject to downstream processing. Read pairs were aligned to the *L. sativa* cv. Salinas RefSeq genome assembly version 7 (genome ID: 5962908) or *S. lycopersicum* cv. Heinz 1706 RefSeq genome assembly (RefSeq GCF_000188115.4) using HISAT2 (v. 2.2.1) with default parameter settings (60). Genes with multiple copies undifferentiated in the genome annotation were assigned numbers in the order they are referred to in the text (e.g., LOC111908039 as *HY5-1*). Mapped reads were assigned to genomic features based on Lsat_Salinas_v7 or *S. lycopersicum* RefSeq assembly annotations using featureCounts (v. 2.0.1) (61). Read counts were summarized at the gene level and zero-count genes were removed prior to further analysis. Raw data and counts have been deposited in NCBI’s Gene Expression Omnibus (62) and are accessible through GEO Series accession numbers GSE180179 and GSE200978 (https://www.ncbi.nlm.nih.gov/geo/query/acc.cgi?acc=GSE180179; https://www.ncbi.nlm.nih.gov/geo/query/acc.cgi?acc=GSE200978).

Differential expression analysis was performed independently for each species in R using the edgeR package (v. 3.34.0) (63, 64). The estimateGLMCommonDisp function was used to estimate a common gene-wise dispersion parameter suitable for all genes and evaluated on an individual basis and likelihood ratio tests were performed to test for differential expression of genes within pairwise treatment groups. For each test, a single treatment (OSC filter) group was compared to the control (clear or shaded glass) treatment and significance was evaluated based on the Benjamini Hochberg adjusted p-value (threshold of FDR<0.05). A second round of analysis was performed by comparing each treatment with the corresponding treatment with variable light intensity.

### Network analysis

Normalized read counts were extracted using the edgeR cpm function and log-transformed counts per million were used as input for weighted correlation network analysis (WGCNA) (v. 1.69) (63, 65, 66). To reduce spurious correlations, genes with consistently low expression (less than two read counts) for four or more samples were removed prior to analysis. With the remaining genes, expression similarity was calculated using the Pearson correlation metric, and a signed adjacency matrix was constructed using a soft-threshold power of 7 for the lettuce dataset and 14 for the tomato dataset to satisfy the scale-free network topology criterion. The network adjacency matrix was then used to calculate the topological overlap for each of the datasets separately. Average linkage hierarchical clustering was performed on the topological overlap dissimilarity matrices, and modules were detected using the dynamic tree cutting algorithm. The resulting network for each species was compared to networks constructed with transcriptome data where both light intensity and quality varied for each species. Topological overlap measures were used to scale these light intensity-dependent networks to the networks with consistent light intensity prior to consensus module detection.

### Statistical analysis

Physiological data were analyzed by ANOVA and Tukey post-hoc test where p<0.05. Treatments that share a letter were not significantly different. Physiological data reported in the body of this report were collected from the same replication of the experiments used in the transcriptome analysis. Photosynthetic data were further analyzed by dividing by the PPFD recorded at time of measurement to identify potential effects of small changes in light intensity introduced by filter degradation. To minimize variation between lettuce replications, the biomass and secondary data were normalized relative to the control treatment within each round. Lettuce biomass and secondary metabolite data were normalized by replicate relative to their respective controls. Two replicates were performed simultaneously in the same growth chamber and were normalized together. These normalized data are presented in the supplementary information.

## Supporting information

DataS1

DataS2

DataS3

DataS4

## Acknowledgments

We would like to thank Jennifer Swift for technical assistance as well as the NCSU Phytotron and the Hernandez group for equipment and advice.

## Author contributions

Conceptualization: CS, WY, HA, BO, HS

Investigation: MC, ER, JC, RH, JR

Software: MC, BE

Visualization: MC, BE, HS

Supervision: CS, WY, HA, BO, HS

Writing—original draft: MC, HS, BE

Writing—review & editing: MC, BE, HA, BO, HS

## Data and materials availability

Raw and processed (count) files from the transcriptome analysis are available in NCBI’s Gene Expression Omnibus and are accessible through GEO Series accession numbers GSE180179 (https://www.ncbi.nlm.nih.gov/geo/query/acc.cgi?acc=GSE180179) and GSE200978 (https://www.ncbi.nlm.nih.gov/geo/query/acc.cgi?acc=GSE200978). All software programs used are open access and freely available.

