## Supplemental Figures & Tables for "Beyond energy balance in agrivoltaic food production: Emergent crop traits from wavelength-selective solar cells"

**This PDF file includes:**

Figs. S1 to S13
Tables S1 to S3
Data S1 to S4

**Other Supplementary Materials for this manuscript include the following:**
Data S1 to S4

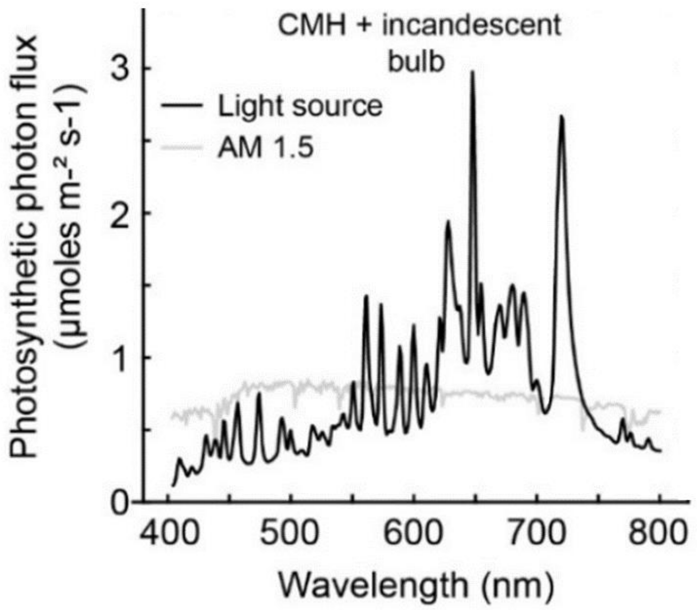


**Fig. S1. Light spectrum in growth chamber**

Light spectrum created with ceramic metal halide (CMH) and incandescent light bulbs in the growth chamber to approximate the spectrum of natural sunlight (AM 1.5) (adapted from Reference 33).

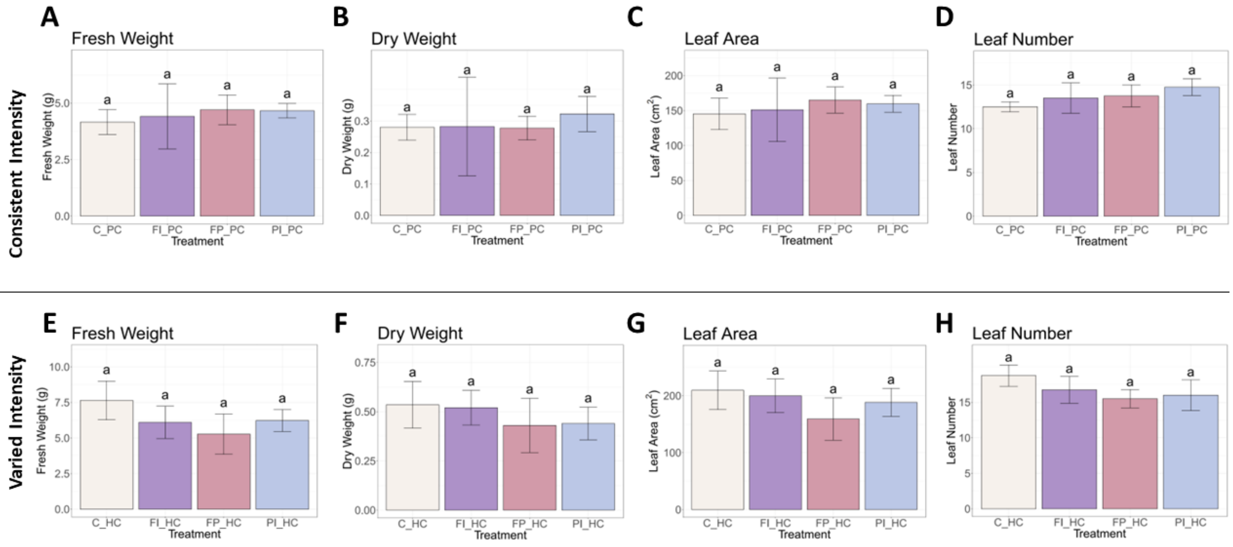


**Fig. S2. Biomass accumulation at 21-day transplant stage in lettuce**

Biomass of the same replication used in the transcriptome analysis at the early harvest, 21-day transplant stage in lettuce under consistent (PC) and varied (HC) light intensity. **A**) Fresh weight of lettuce in the PC experiment. **B**) Dry weight in the PC experiment. **C**) Leaf area in the PC experiment. **D**) Total leaf number per plant in the PC experiment. **E-H**) Fresh weight, dry weight, leaf area and leaf number in the HC experiment (adapted from Reference 33). Error bars represent the standard deviation. Statistical significance was assessed by ANOVA and Tukey test ($p<0.05$).

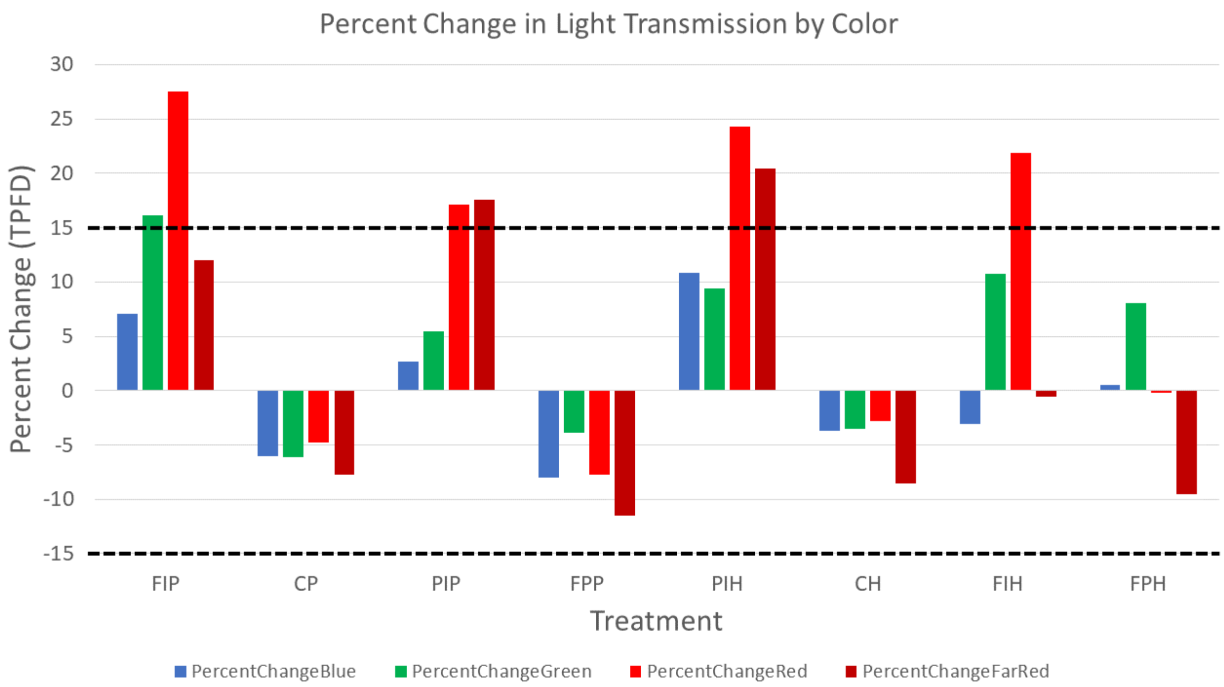


**Fig. S3. Percent change in filter transmission by color in lettuce experiments**

Percent change in PFD for blue (400-500 nm), green (500-600 nm), red (600-700) and far-red (700-750 nm) light measured before and after each replication. Both treatments where light intensity was consistent (**P) and varied (**H) were considered. Dashed horizontal lines represent the 15% percent change that was considered to be due to actual filter degradation and not instrument or lighting variation. Filter degradation was observed by the end of each replication. This resulted in a small shift in spectral composition and increase in light intensity reaching the plants. Most of the percent change in blue, green, red and far-red photon flux density was less than 15% for both experiments. Variations less than 15% were also seen in the clear and shaded glass controls and were attributed to instrument error or variation in the light source.

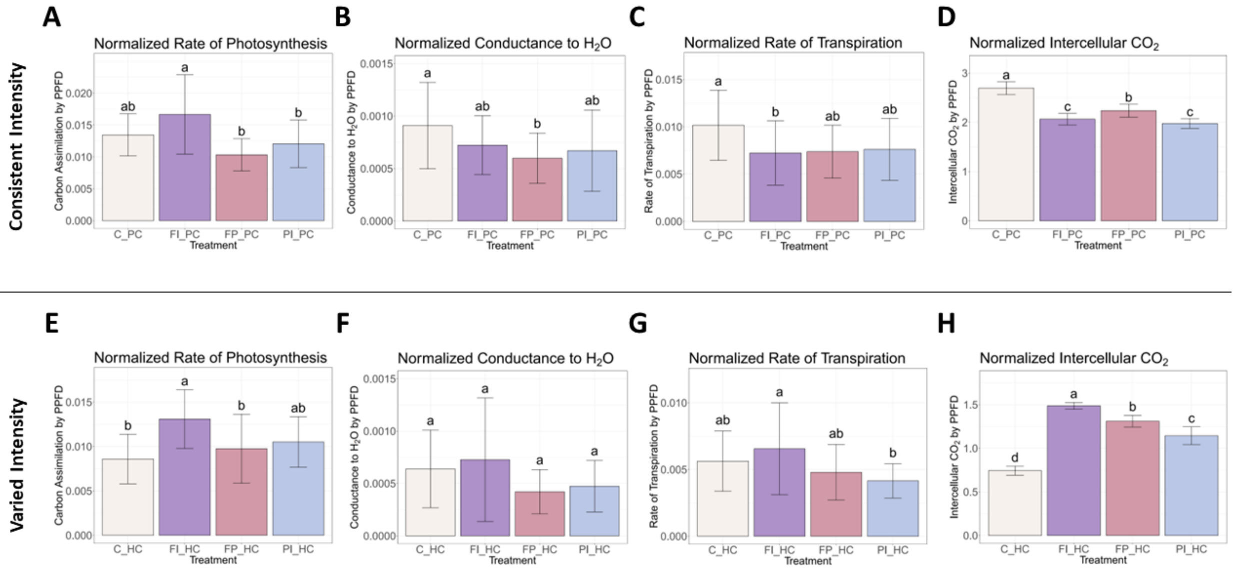


**Fig. S4. Lettuce photosynthetic measurements normalized to incident PPFD**
Carbon assimilation and other measurements normalized to the PPFD recorded by the instrument at the time the measurements were taken in lettuce experiments under consistent (PC) and varied (HC) light intensity. **A**) Normalized rate of photosynthesis measured under each light treatment of the PC experiment. **B**) Normalized stomatal conductance to water. **C**) Normalized transpiration rate. **D**) Normalized intercellular $CO_2$. **E-H**) Normalized measurements of the HC experiment (adapted from Reference 33). Error bars represent the standard deviation. Statistical significance was assessed by ANOVA and Tukey test ($p<0.05$).

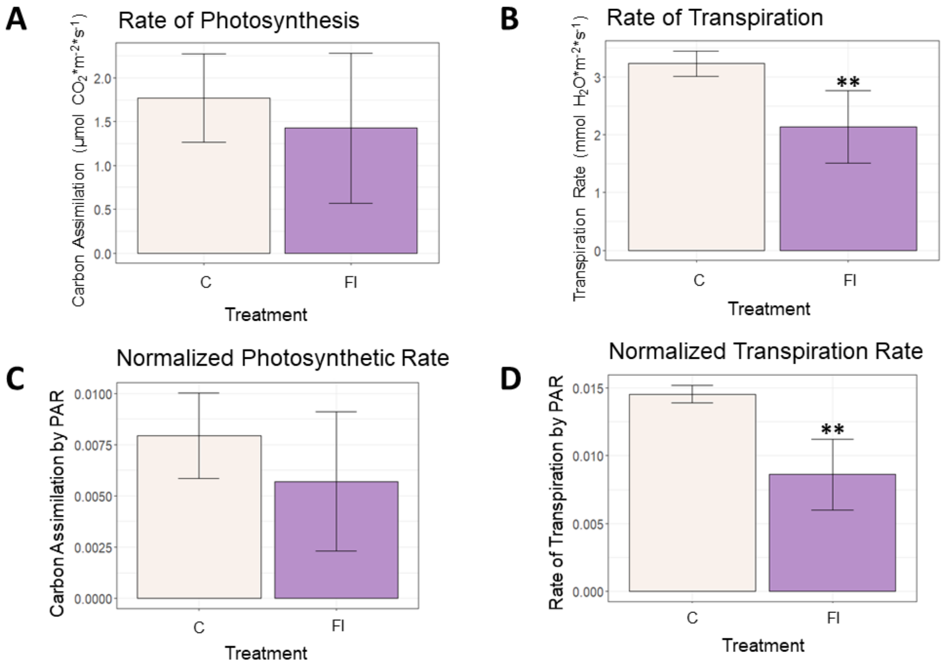


**Fig. S5. Tomato photosynthetic measurements under consistent light intensity**
**A**) Rate of photosynthesis measured under the FI filter and a spectrally neutral control with the same light intensity. **B**) Rate of transpiration under the FI filter and control. **C**) Rate of photosynthesis under the FI filter and control normalized by incident PPFD measured by the Li-cor during each measurement. **D**) Normalized rate of transpiration under the FI filter and control. Small variation in PPFD at the time of measurement had no meaningful effect. Significance levels indicated where relevant (*$p<0.05$,**$p<0.01$,***$p<0.001$).

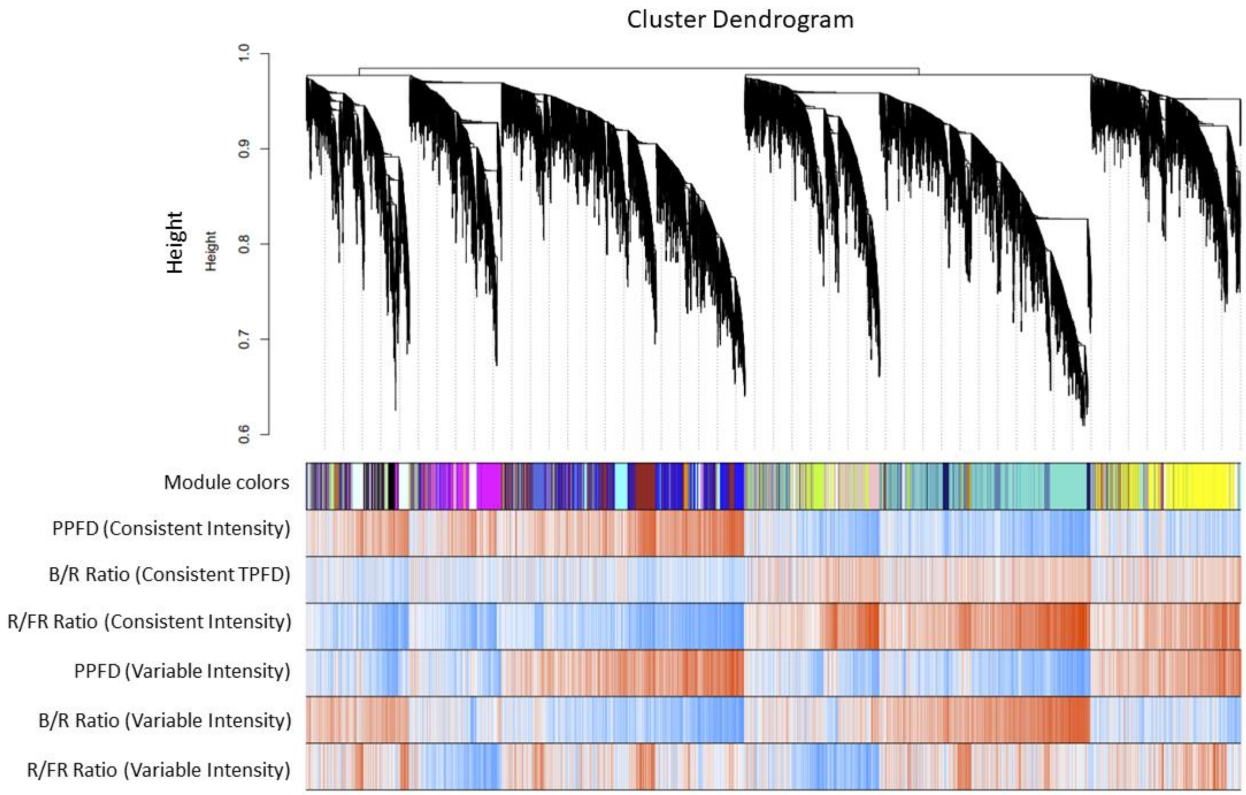


**Fig. S6. Dendrogram of lettuce gene expression under consistent and varied light intensity** Dendrogram of gene modules from signed WGCNA from lettuce leaf tissue. More closely related modules are separated by fewer branches. Red and blue coloring indicates correlation between modules and different light conditions under consistent and variable light intensity. Genes that were more highly expressed when PPFD increased tended to have lower expression when B/R or R/FR ratios increased.

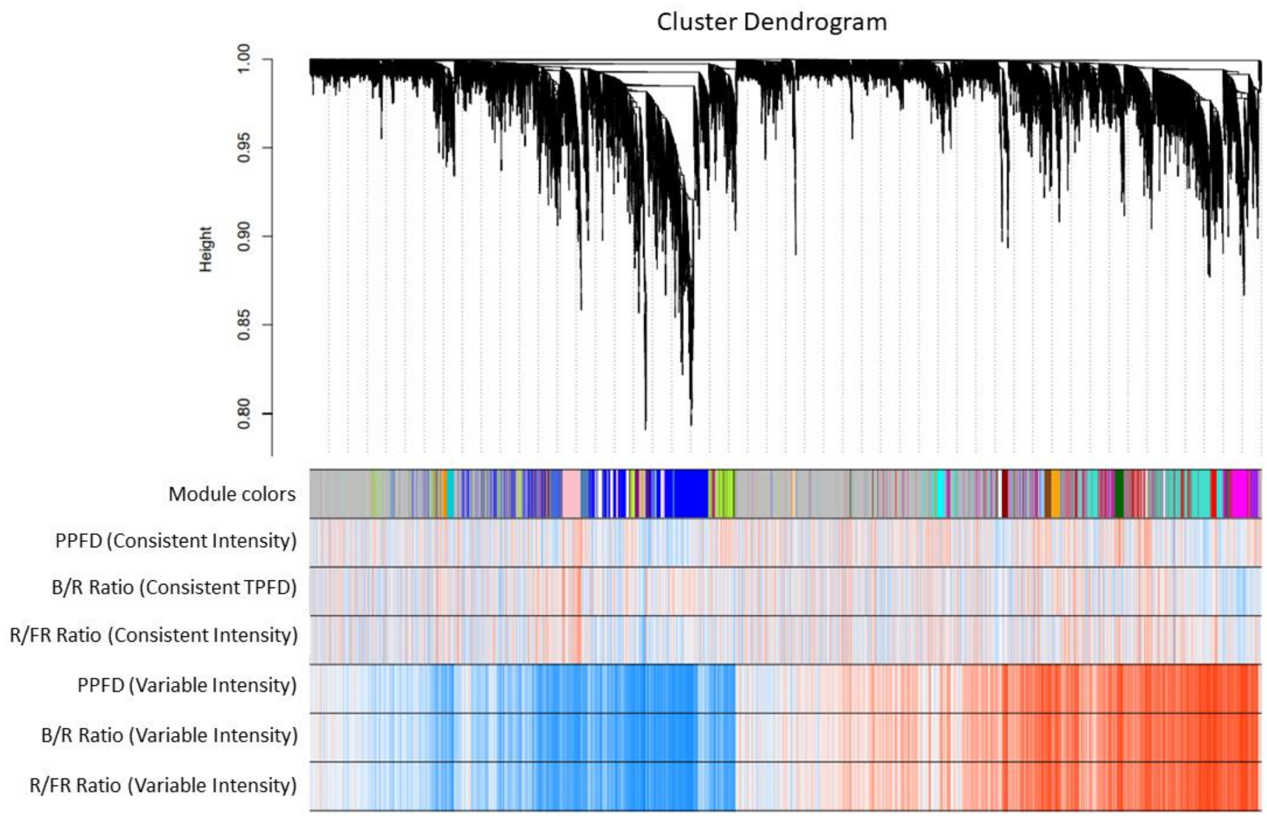


**Fig. S7. Dendrogram of tomato gene expression under consistent and varied light intensity**
Dendrogram of gene modules from signed WGCNA from tomato leaf tissue. More closely related modules are separated by fewer branches. Red and blue coloring indicates correlation between modules and different light conditions under consistent and variable light intensity. Stronger correlations between gene expression and light traits were observed when light intensity was varied.

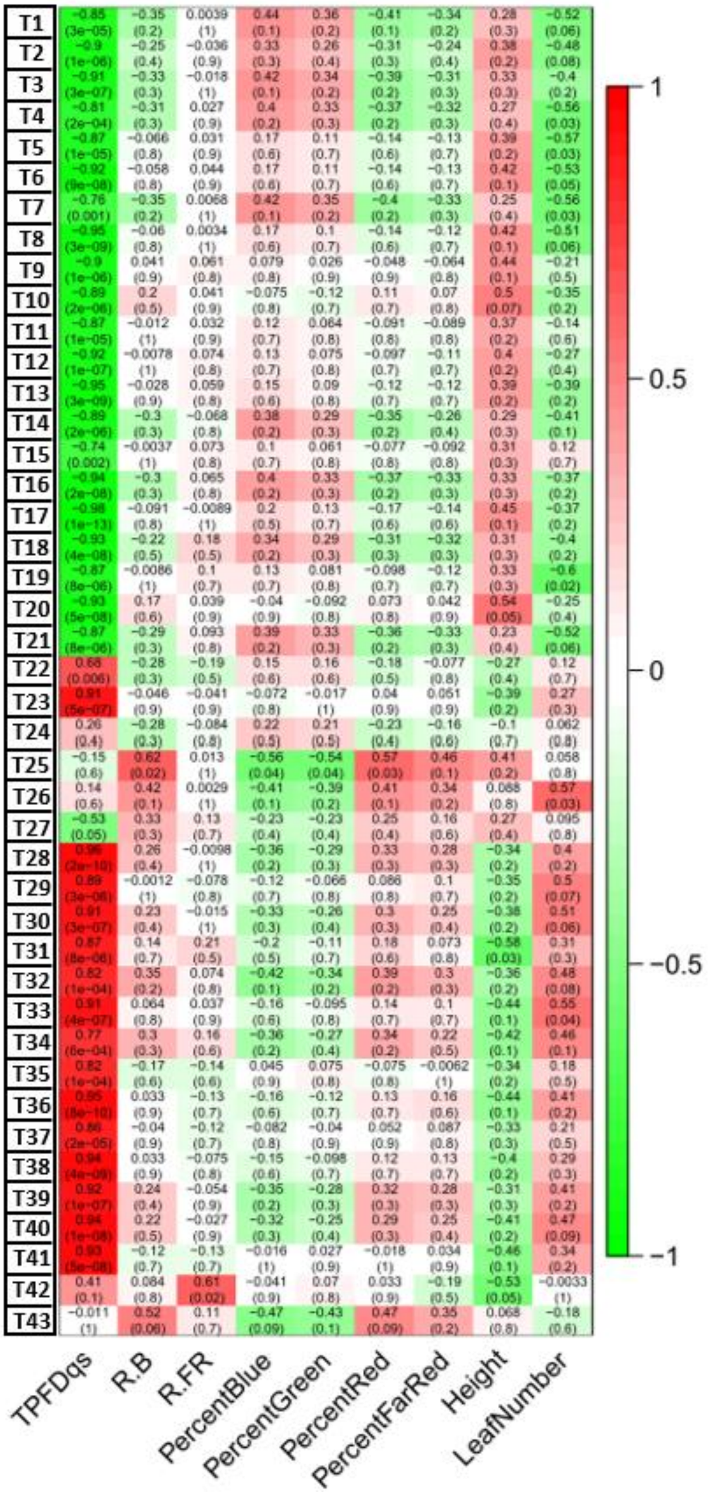


**Fig. S8. Module-trait relationships in tomato genes under varied light intensity**
Module-trait relationships of gene modules from signed WGCNA from tomato leaf tissue grown under variable light intensity. Significant correlations between the module eigengenes of light and physiological traits are indicated with either red (positive) and green (negative). Many modules significantly correlated with TPFD and significant correlations with all other traits except percent far-red light.

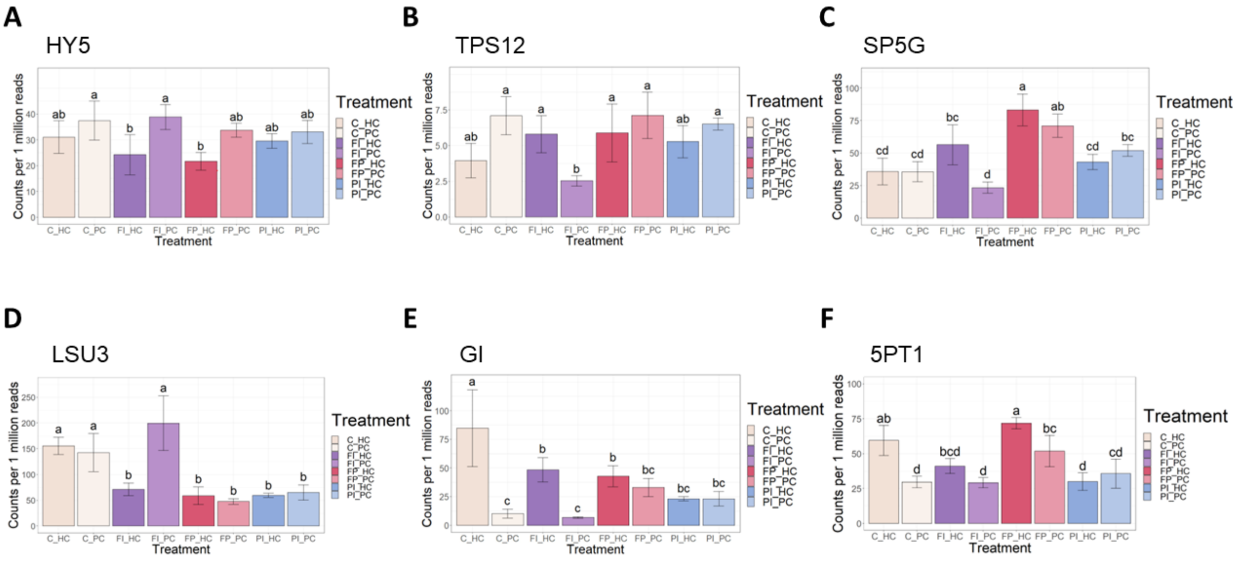


**Fig. S9. Expression levels of select genes of interest in tomato**
Normalized expression of genes identified in the tomato genome. **A**) Expression of *elongated hypocotyl 5* (*HY5*) in tomato leaves under different filters with consistent and variable light intensity, measured in counts per million (CPM). **B**) Expression of *terpene synthase 12* (*TPS12*). **C**) Expression of *self pruning 5G* (*SP5G*). **D**) Expression of *response to low sulfur 3-like* (*LSU3*). **E**) Expression of *gigantea* (*GI*). **F**) Expression of *inositol-1,4,5-triphosphate-5-phosphatase* (*5PT1*). Error bars represent the standard deviation. Letters indicate significance from ANOVA and Tukey test ($p<0.05$).

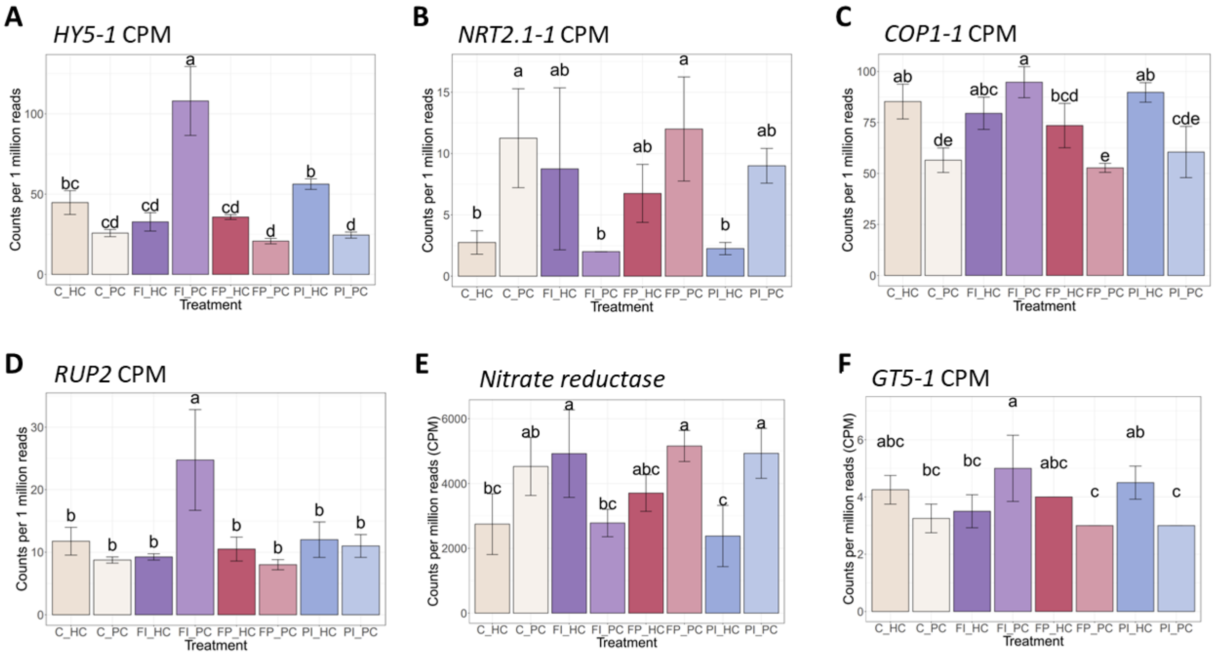


**Fig. S10. Expression levels of select genes of interest in lettuce**

Normalized expression of genes identified in the lettuce genome, measured in counts per million (CPM). **A**) Expression of *elongated hypocotyl 5* (*HY5*) in lettuce leaves under different filters with consistent and variable light intensity. **B**) Expression of *high-affinity nitrate transporter 2.1* (*NRT2.1-1*). **C**) Expression of *constitutive photomorphogenic 1* (*COP1*). **D**) Expression of *repressor of UV-B photomorphogenesis 2* (*RUP2*). **E**) Expression of *nitrate reductase* (*NR*). **F**) Expression of *anthocyanidin 3-O-glucosyltransferase 5* (*GT5*). Error bars represent the standard deviation. Letters indicate significance from ANOVA and Tukey test ($p<0.05$).

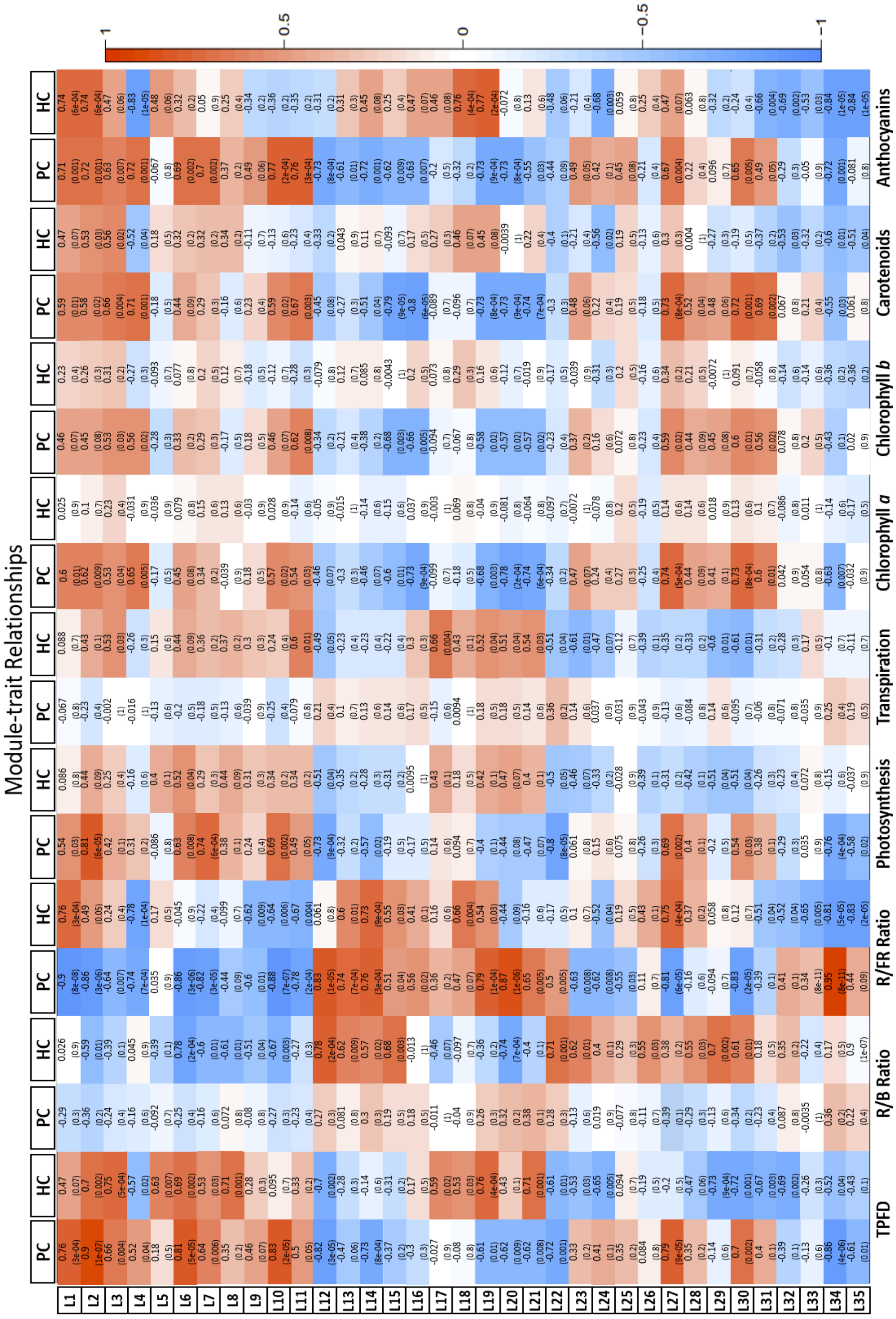


**Fig. S11. Module-trait relationships of lettuce**

Table of correlations between measured light and physiological parameters and the 35 modules identified in the network analysis of lettuce under consistent (PC) and varied (HC) light intensity. Strong positive correlations are dark red while strong negative correlations are dark blue. Values given are the correlation coefficient (*r*) for each relationship. Correlations with p<1e-05 are significant, given in parentheses below *r*.

Module-trait Relationships

| Module | TPFD | R/B Ratio | R/FR Ratio | Percent Blue | Percent Green | Percent Red | Percent Far-red | Height | Leaf Number |
|---|---|---|---|---|---|---|---|---|---|
| T1 | -0.097 (0.7) | | 0.0039 (1) | | 0.0058 (1) | | -0.058 (0.8) | | |
| T2 | -0.14 (0.6) | | | | | | | | |
| T3 | | | | | | | | 0.23 (0.4) | -0.14 (0.6) |
| T4 | -0.12 (0.7) | | | | | | | 0.17 (0.5) | -0.18 (0.5) |
| T5 | | | | | | | | 0.35 (0.2) | |
| T6 | | -0.0086 (1) | 0.044 (0.9) | 0.075 (0.8) | 0.069 (0.8) | -0.049 (0.9) | -0.13 (0.7) | 0.21 (0.4) | |
| T7 | -0.18 (0.5) | -0.2 (0.5) | 0.0068 (1) | 0.24 (0.4) | 0.29 (0.3) | -0.25 (0.4) | -0.32 (0.3) | | -0.11 (0.7) |
| T8 | -0.31 (0.2) | | 0.0034 (1) | | 0.066 (0.8) | | -0.12 (0.7) | 0.28 (0.3) | -0.14 (0.6) |
| T9 | | 0.041 (0.9) | 0.061 (0.8) | | 0.016 (1) | | -0.064 (0.8) | 0.095 (0.7) | |
| T10 | -0.026 (0.9) | 0.2 (0.5) | 0.041 (0.9) | -0.075 (0.8) | | 0.11 (0.7) | | | |
| T11 | -0.11 (0.7) | | 0.032 (0.9) | | | | | | -0.14 (0.6) |
| T12 | | | 0.074 (0.8) | | | | | 0.043 (0.9) | -0.14 (0.6) |
| T13 | -0.062 (0.8) | | 0.059 (0.8) | | | | | 0.31 (0.2) | -0.39 (0.2) |
| T14 | | | -0.068 (0.8) | | | | | 0.29 (0.3) | |
| T15 | | -0.0037 (1) | | 0.017 (1) | | -0.01 (1) | | | |
| T16 | -0.092 (0.7) | -0.14 (0.6) | | 0.04 (0.9) | | -0.039 (0.9) | | 0.25 (0.4) | -0.25 (0.4) |
| T17 | -0.13 (0.6) | | -0.0089 (1) | | | | | 0.14 (0.6) | -0.18 (0.5) |
| T18 | | | | | | | | 0.077 (0.8) | |
| T19 | | | | | | | | 0.22 (0.4) | -0.16 (0.6) |
| T20 | | | | -0.0067 (1) | -0.092 (0.8) | 0.029 (0.9) | 0.042 (0.9) | 0.49 (0.05) | |
| T21 | | -0.17 (0.5) | | 0.14 (0.6) | 0.052 (0.9) | -0.12 (0.7) | -0.014 (1) | | -0.28 (0.3) |
| T22 | 0.096 (0.7) | -0.28 (0.3) | -0.19 (0.5) | 0.15 (0.6) | 0.16 (0.6) | -0.18 (0.5) | -0.077 (0.8) | | 0.12 (0.7) |
| T23 | | -0.046 (0.9) | -0.041 (0.9) | | | | | -0.15 (0.6) | |
| T24 | | -0.16 (0.6) | -0.084 (0.8) | 0.13 (0.6) | 0.092 (0.7) | -0.13 (0.6) | -0.051 (0.9) | -0.1 (0.7) | |
| T25 | | | | | | | 0.025 (0.9) | 0.41 (0.2) | 0.058 (0.8) |
| T26 | 0.067 (0.8) | | | | | | | | 0.075 (0.8) |
| T27 | | 0.26 (0.3) | 0.11 (0.7) | -0.2 (0.5) | -0.18 (0.5) | 0.22 (0.4) | 0.12 (0.7) | | 0.095 (0.8) |
| T28 | 0.25 (0.3) | | -0.0098 (1) | -0.024 (0.9) | -0.16 (0.6) | 0.059 (0.8) | 0.2 (0.5) | | 0.11 (0.7) |
| T29 | | -0.0012 (1) | -0.078 (0.8) | | | | | -0.073 (0.8) | |
| T30 | 0.12 (0.7) | | -0.015 (1) | | | | | -0.11 (0.7) | 0.077 (0.8) |
| T31 | 0.07 (0.8) | 0.14 (0.7) | | -0.2 (0.5) | -0.11 (0.7) | 0.18 (0.6) | 0.073 (0.8) | | |
| T32 | 0.41 (0.1) | 0.34 (0.2) | 0.0062 (1) | -0.28 (0.3) | -0.28 (0.3) | 0.31 (0.3) | 0.22 (0.4) | -0.065 (0.8) | 0.13 (0.6) |
| T33 | 0.24 (0.4) | 0.064 (0.8) | 0.037 (0.9) | -0.16 (0.6) | -0.095 (0.8) | 0.14 (0.7) | 0.1 (0.7) | -0.0082 (1) | |
| T34 | 0.11 (0.7) | 0.3 (0.3) | 0.12 (0.7) | -0.36 (0.2) | -0.27 (0.4) | 0.34 (0.2) | 0.22 (0.5) | | |
| T35 | | -0.17 (0.6) | -0.0027 (1) | 0.045 (0.9) | 0.075 (0.8) | -0.075 (0.8) | -0.0062 (1) | -0.1 (0.7) | 0.18 (0.5) |
| T36 | | | | | | | | -0.44 (0.1) | |
| T37 | | -0.04 (0.9) | -0.028 (0.9) | | | | | -0.037 (0.9) | 0.16 (0.6) |
| T38 | 0.12 (0.7) | | | | | | | -0.2 (0.5) | 0.19 (0.5) |
| T39 | 0.086 (0.8) | | | | | | | | 0.41 (0.2) |
| T40 | 0.21 (0.4) | 0.22 (0.5) | | -0.28 (0.3) | -0.14 (0.6) | 0.28 (0.3) | 0.014 (1) | -0.18 (0.5) | 0.11 (0.7) |
| T41 | 0.055 (0.8) | | | -0.016 (1) | | | 0.034 (0.9) | -0.46 (0.1) | |
| T42 | 0.25 (0.4) | 0.084 (0.8) | 0.0023 (1) | -0.041 (0.9) | | 0.033 (0.9) | | -0.45 (0.08) | |
| T43 | | 0.22 (0.4) | 0.11 (0.7) | -0.14 (0.6) | -0.0093 (1) | 0.12 (0.7) | | 0.025 (0.9) | |

**Fig. S12. Module-trait relationships of tomato**

Table of correlations between measured light and physiological parameters and the 43 modules identified in the network analysis of tomato under consistent (PC) and varied (HC) light intensity. Strong positive correlations are dark red while strong negative correlations are dark green. Values given are the correlation coefficient (*r*) for each relationship. Non-significant correlations are indicated in gray. Correlations with p<1e-05 are significant, given in parentheses below *r*.

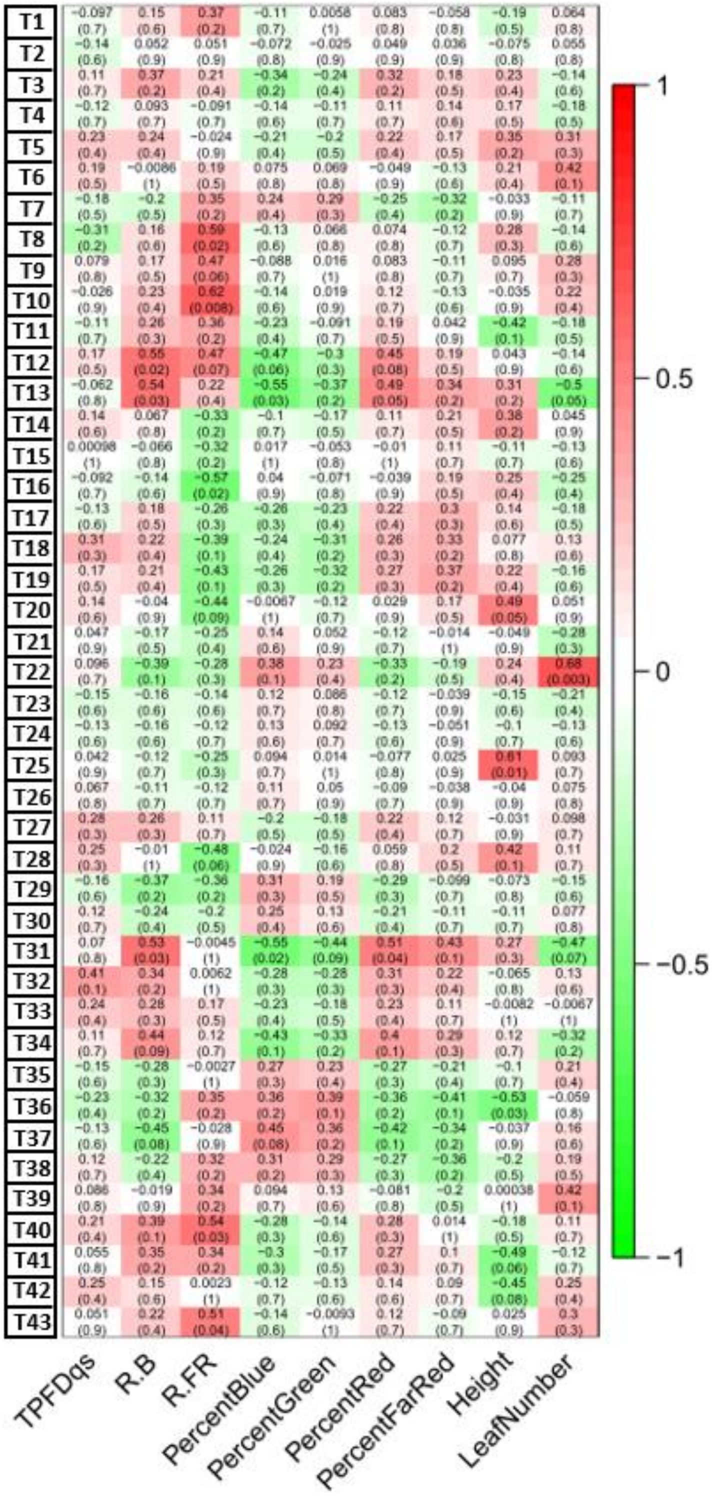


**Fig. S13. Module-trait relationships in tomato genes under consistent light intensity**
Module-trait relationships of gene modules from signed WGCNA from tomato leaf tissue grown under variable light intensity. Significant correlations between the module eigengenes of light and physiological traits are indicated with either red (positive) and green (negative). Unlike in lettuce, there were no modules significantly correlated with TPFD. There were significant correlations with all other traits except percent green light.

| Gene | Product | FDR | $log_2FC$ |
|---|---|---|---|
| LOC111902387 | transcription factor MYB114 | 2.23E-10 | 1.5 |
| LOC111908039 | transcription factor HY5-1 | 1.24E-49 | 2.2 |
| LOC111883156 | transcription factor HY5-2 | 2.51E-17 | 1.5 |
| LOC111918210 | transcription factor HHO5 | 3.24E-27 | -1.1 |
| LOC111884885 | zinc finger protein CONSTANS-LIKE 5-1 | 1.60E-25 | 1.3 |
| LOC111880753 | suppressor of overexpression of CO 1 | 5.86E-01 | 1.6 |
| LOC111897754 | high affinity nitrate transporter 2.4-1 | 5.11E-34 | -2.8 |
| LOC111886104 | high affinity nitrate transporter 2.1-1 | 2.33E-27 | -6.8 |
| LOC111886119 | high affinity nitrate transporter 2.1-5 | 1.86E-08 | -3.7 |
| LOC111897821 | high affinity nitrate transporter 2.4-2 | 5.32E-07 | -3.0 |
| LOC111910637 | high affinity nitrate transporter 2.5 | 6.98E-06 | -2.0 |
| LOC111886074 | high affinity nitrate transporter 2.1-2 | 1.04E-07 | -5.5 |
| LOC111876411 | high affinity nitrate transporter 2.7 | 1.00E-02 | 1.3 |
| LOC111886105 | high affinity nitrate transporter 2.1-4 | 4.07E-02 | -3.3 |
| LOC111893654 | nitrate reductase [NADH] | 1.83E-05 | -0.7 |
| LOC111902386 | transcription factor MYB75 (PAP1) | 9.95E-13 | 1.1 |
| LOC111883451 | chalcone synthase | 9.18E-01 | 0.3 |
| LOC111900153 | flavonol synthase/flavanone 3-hydroxylase | 6.52E-01 | 0.4 |
| LOC111911650 | anthocyanidin 3-O-glucosyltransferase 2-1 | 1.05E-10 | 2.4 |
| LOC111911648 | anthocyanidin 3-O-glucosyltransferase 2-2 | 2.94E-03 | 1.0 |
| LOC111906208 | anthocyanin 3-O-glucoside-6''-O-malonyltransferase | 1.30E-04 | 0.4 |
| LOC111897116 | (3S,6E)-nerolidol synthase 1-1 | 2.02E-05 | 2.7 |

**Table S1. Selected DEGs from lettuce FI/C comparison**

$Log_2$ fold changes and adjusted p-values (FDRs) are given for selected genes differentially expressed (DEGs) between the FI filter treatment and the shaded glass control in lettuce with consistent light intensity. Both *HY5* genes and several other transcription factors are significantly up- or downregulated. Other genes involved in nitrate transport and flowering are also differentially expressed. Genes involved in late anthocyanin biosynthesis and modification, but not early biosynthesis, are upregulated. $Log_2$ fold changes with $FDR<0.05$ are considered significant.

| Treatment Contrast | Genes | logFC | logCPM | LR | P-value | FDR | Product |
|---|---|---|---|---|---|---|---|
| FP_PC | LOC101243970 | -1.58666 | 6.622486 | 56.27023 | 6.32E-14 | 1.30E-09 | protein RESPONSE TO LOW SULFUR 3-like |
| FP_PC | LOC101255788 | 1.684499 | 5.011209 | 52.33528 | 4.68E-13 | 4.80E-09 | protein GIGANTEA |
| FP_PC | LOC101252466 | -1.57571 | 3.468235 | 29.60999 | 5.28E-08 | 0.000298 | 5'-adenylylsulfate reductase 3%2C chloroplastic-like |
| FP_PC | LOC101257319 | -1.41301 | 5.951838 | 29.42663 | 5.81E-08 | 0.000298 | 5'-adenylylsulfate reductase 3%2C chloroplastic |
| FP_PC | SP5G | 0.985859 | 5.672479 | 27.91465 | 1.27E-07 | 0.000521 | protein SELF PRUNING 5G |
| FP_PC | LOC101254785 | 0.773774 | 6.830249 | 25.27077 | 4.98E-07 | 0.001705 | two-component response regulator-like PRR37 |
| FP_PC | LOC101246670 | 1.300331 | 5.665381 | 23.99726 | 9.65E-07 | 0.00283 | ABC transporter G family member 11-like |
| FP_PC | LOC101268293 | 0.811524 | 4.611487 | 23.11674 | 1.52E-06 | 0.003913 | auxin-responsive protein SAUR78 |
| FP_PC | 5PT1 | 0.791617 | 5.437434 | 21.46818 | 3.60E-06 | 0.008165 | inositol-1,4,5-triphosphate-5-phosphatase |
| FP_PC | LOC101259102 | 2.822703 | -1.24142 | 21.27605 | 3.98E-06 | 0.008165 | ras GTPase-activating protein-binding protein 2-like |
| FP_PC | LOC544267 | -0.93581 | 9.482485 | 20.49607 | 5.98E-06 | 0.011153 | adenylyl-sulfate reductase |
| FP_PC | GLK2 | 0.786459 | 4.537453 | 19.96566 | 7.88E-06 | 0.01269 | golden2-like protein |
| FP_PC | LOC101251881 | -0.7291 | 4.808613 | 19.87068 | 8.29E-06 | 0.01269 | protein LTV1 homolog |
| FP_PC | LOC101266832 | -0.60265 | 4.739285 | 19.78779 | 8.65E-06 | 0.01269 | uncharacterized LOC101266832 |
| PI_PC | LOC101243970 | -1.14345 | 6.622486 | 29.9733 | 4.38E-08 | 0.000899 | protein RESPONSE TO LOW SULFUR 3-like |
| PI_PC | LOC101255788 | 1.17469 | 5.011209 | 25.78052 | 3.83E-07 | 0.003927 | protein GIGANTEA |
| PI_PC | LOC101246103 | -1.94398 | -0.10848 | 23.02473 | 1.60E-06 | 0.010945 | zingipain-2-like |
| FI_PC | LOC104648118 | -1.48387 | 2.519074 | 33.00988 | 9.17E-09 | 0.000188 | (E%2CE)-germacrene B synthase-like |
| FI_PC | LOC101250428 | -1.04147 | 3.442472 | 21.87674 | 2.91E-06 | 0.029846 | terpene synthase |
| FI_PC | MKS1d | -1.01486 | 4.323152 | 21.03354 | 4.51E-06 | 0.030886 | methylketone synthase Id |
| FI_PC | LOC101245161 | -2.99589 | 1.048603 | 20.34504 | 6.47E-06 | 0.033189 | GeneID:101245161 |

**Table S2. Selected DEGs from tomato filter/C comparison**

$Log_2$ fold changes and adjusted p-values (FDRs) are given for selected genes differentially expressed (DEGs) between the FP, PI and FI filter treatments and the shaded glass control in tomato with consistent light intensity. The majority of the differentially expressed genes were found in the FP/C contrast. $Log_2$ fold changes with FDR<0.05 are considered significant.

| Treatment | Lettuce | | | | Tomato | | | |
|---|---|---|---|---|---|---|---|---|
| Blue (400-500nm) (% of TFPD) | 107 (14%) | 60 (14%) | 53 (12%) | 78 (15%) | 84 (14%) | 44 (16%) | 33 (11%) | 49 (19%) |
| Green (500-600nm) (% of TFPD) | 223 (30%) | 111 (27%) | 119 (26%) | 158 (31%) | 173 (30%) | 84 (30%) | 75 (26%) | 90 (35%) |
| Red (600-700nm) (% of TFPD) | 353 (47%) | 205 (50%) | 243 (53%) | 236 (46%) | 277 (48%) | 130 (46%) | 153 (53%) | 103 (40%) |
| Far-red (700-750nm) (% of TFPD) | 60 (8%) | 37 (9%) | 41 (9%) | 39 (8%) | 106 (8%) | 56 (9%) | 59 (9%) | 36 (6%) |
| PPFD (400-700nm) | 684 | 376 | 415 | 473 | 581 | 282 | 288 | 258 |
| TFPD (400-750 nm) | 743 | 413 | 457 | 511 | 687 | 338 | 347 | 294 |

**Table S3. Light intensity and spectra in variable light OSC filter treatments**
Breakdown of the photon flux (μmol $m^{-2}$ $s^{-1}$) reaching the plants by color measured at the end of the experiments. The percentages of each color relative to the total photon flux density (TPFD) were consistent for each filter in both experiments. TPFD variation in lettuce was more distributed than in tomato, where TPFD of the filter treatments varied only by 53 μmol $m^{-2}$ $s^{-1}$.

**Data S1. Light conditions by treatment**

Characteristics of the light conditions in each treatment used in the weighted gene correlation network analyses (WGCNA) of lettuce and tomato. Light conditions were measured in several points in each growth both and were averaged to a single value for each treatment. Amounts of red, blue, etc. are included as well as R/FR and R/B ratios. (See Excel file.)

**Data S2. Physiological data**

Raw biomass and carbon assimilation data used to generate lettuce and tomato figures. Secondary metabolite data from lettuce and flowering dates from tomato. Means and standard deviations are shown. Significant differences are indicated with ANOVA p-values and results from Tukey tests where applicable. (See Excel file.)

**Data S3. Compiled DEGs from differential gene analyses**

Differentially expressed genes (DEGs) from comparisons of each OSC filter to the relative control for consistent (PC) and variable light intensity (HC) experiments as well as DEGs from HC/PC comparisons for each filter and control spectrum. (See Excel file.)

**Data S4. Module assignments of lettuce and tomato genes**

Weighted gene correlation network analyses (WGCNA) for lettuce and tomato experiments. The consistent light intensity experiment (PC) data are referred to as "Set 1" while the variable light intensity experiment (HC) is "Set 2". The module numbers used in the text are given under "ModuleNumber". "ModuleLabel" and "ModuleColor" are auto-assigned labels that were not used in the text. (See Excel file.)
